# Bacterial inoculation elicits changes to the coral epigenome

**DOI:** 10.1101/2024.10.27.620496

**Authors:** Adam R. Barno, Helena D.M. Villela, Pedro M. Cardoso, Francisca C. García, Guoxin Cui, Nathalia Delgadillo-Ordoñez, Alexandre S. Rosado, Torsten Thomas, Manuel Aranda, Christian R. Voolstra, Raquel S. Peixoto

## Abstract

Environmental shifts can cause epigenetic modifications in corals, which are associated with changes in gene expression and physiology, though it remains unclear if associated bacteria can also induce such changes. Here, we inoculated nubbins of the coral *Pocillopora verrucosa* with an opportunistic pathogen, *Vibrio coralliilyticus*, and/or a coral probiotic, *Cobetia* sp., and subjected the nubbins to heat stress. We show that pathogen exposure led to distinct DNA methylation changes compared to the control, probiotic, and co-inoculation groups. We also demonstrate that DNA methylation correlates with coral gene expression and highlight genes altered by pathogen inoculation that showed similar responses in their expression and methylation. Notably, the coral probiotic was able to mitigate specific epigenetic changes, which correlated with increased stress resilience and higher coral survival rates. Thus, bacterial-induced changes to the coral epigenome may instigate long-term changes in host resilience.

## Main

Recent work has shown that beneficial bacteria (probiotics) can increase coral health and resilience in response to stress caused by pathogens, increased temperature, and oil spills^1–7^. This is because bacteria are essential to the proper functioning of the coral holobiont^8–10^, which consists of the coral animal and its associated symbiotic microbes. For example, inoculation of the coral *Pocillopora damicornis* with beneficial microorganisms for corals (BMCs) consisting of bacteria from the genera *Yangia*, *Roseobacter*, *Phytobacter*, and *Salinicola*, correlated with increases in energy reserves and calcification rates^11^. Furthermore, administration of BMCs during and after heat stress increased transcription of genes involved in recovery and stress attenuation and decreased those related to inflammatory and innate immune responses in the coral *Mussismilia hispida*^5^. These changes also correlated with improved coral survival and increased photosynthetic efficiency in the associated symbiotic microalgae (i.e., Symbiodiniaceae)^5^. High temperatures, though, can also lead to dysbiosis and increased establishment of opportunistic pathogens, causing coral bleaching and death^5,12^. For example, inoculation with the temperature-dependent coral pathogen *Vibrio coralliilyticus* brought about a reduction in the skeletal density and abnormal crystal morphology in *P. damicornis*^3^. This could be attenuated by inoculation with a BMC consortium composed of bacteria from the genera *Pseudoalteromonas*, *Cobetia*, and *Halomonas*^3^, showing the protective capacity of BMCs against pathogens as well as temperature stress.

Coral-associated microorganisms have recently been posited to have the ability to alter coral epigenomes, thus modifying phenotypic responses^13^. The most commonly studied epigenome modification in corals is DNA methylation, which, depending on the location of the methylated bases, prevents transcriptional noise and increases fidelity^14–16^. The interaction between coral epigenomes and their symbionts was recently demonstrated by exchanging the Symbiodiniaceae strain in the coral *Montastraea cavernosa* and applying thermal stress, which together produced changes in DNA methylation^17^. Exchanging or manipulating bacterial symbionts may have similar impacts on the coral host, however this has not been studied. These impacts could be either indirect, by changing the nutritional status or altering metabolic capacities^10^, or direct, by producing metabolites that act as cofactors for epigenome-modifying enzymes^18^ or encoding proteins that are homologs of bacteria-produced proteins capable of inducing epigenomic changes in mammals^13^.

Here, we sought to determine whether inoculation with bacteria leads to changes in the coral epigenome that align with improved resilience to thermal stress. To do this, we regularly inoculated *P. verrucosa* nubbins with either a BMC (*Cobetia* sp. strain pBMC4^19^; hereafter “*Cobetia*”), a temperature-dependent pathogen (*V. coralliilyticus* strain BAA-450^20^; hereafter “*Vibrio*”), or both bacteria (hereafter “Both”), and subjected the corals to a temperature stress regime (from 27 °C to 33 °C), followed by a recovery period. Heat-stressed corals are hereafter denoted with “H_”, whereas corals kept at ambient temperature throughout the experiment are denoted with “A_”. The mesocosm experiment encompassed 28 days, with samples taken before any inoculation was applied (T0 = day 0), at the peak of the heat stress (T1 = day 19), and after a recovery period (T2 = day 28). DNA methylation differences at the gene level were analyzed and linked to increases in the abundance of the respective inocula, gene expression levels, and coral phenotypic responses, to show that bacterial inoculation, in combination with temperature stress, can modulate the coral epigenome. Through co-inoculation of a probiotic and pathogen, we could further show that indicators of stress seen in the pathogen-inoculated corals were reduced with the co-inoculated BMC. This pattern could be observed in all components of the holobiont (host phenotype, host epigenome, host gene expression, symbiont photosynthetic efficiency, and microbiome composition) and underscores the potential for probiotic therapy as prevention of epigenomic disturbances caused by pathogenic infection.

## Results

### Probiotic inoculation increases coral resilience

To evaluate the effects of bacterial inoculation and temperature stress on *P*. *verrucosa*, we monitored nubbin health over a 28-day mesocosm experiment using photosynthetic efficiency measurements and nubbin condition/coloration. This showed that *Vibrio* inoculation negatively affected coral resilience to temperature stress during recovery, while the *Cobetia* and co-inoculation groups showed more resilience. The photosynthetic efficiency (*Fv*/*Fm* values measured by Diving-PAM II) was significantly lower in heat-stressed corals than in ambient temperature corals at T1 (*Wilcoxon Rank Sum*: *W* = 256, p = 1.54e-6) and T2 (*Wilcoxon Rank Sum*: *W* = 248, p = 6.64e-6) (Fig. 1A and Supplementary Table 1a). The *Fv/Fm* values were not significantly different between inoculation groups in either the ambient temperature corals (*Kruskal-Wallis*: *X^2^* = 4.87, df = 3, p = 0.182) nor the heat-stressed corals (*Kruskal-Wallis*: *X^2^* = 3.97, df = 3, p = 0.265) at T2, although the mean *Fv/Fm* for H_*Vibrio* was lower than the other heat-stressed groups (0.368 ± 0.245 for H_*Vibrio*, 0.523 ± 0.047 for H_Control, 0.514 ± 0.045 for H_*Cobetia*, 0.530 ± 0.012 for H_Both).

**Fig. 1.**
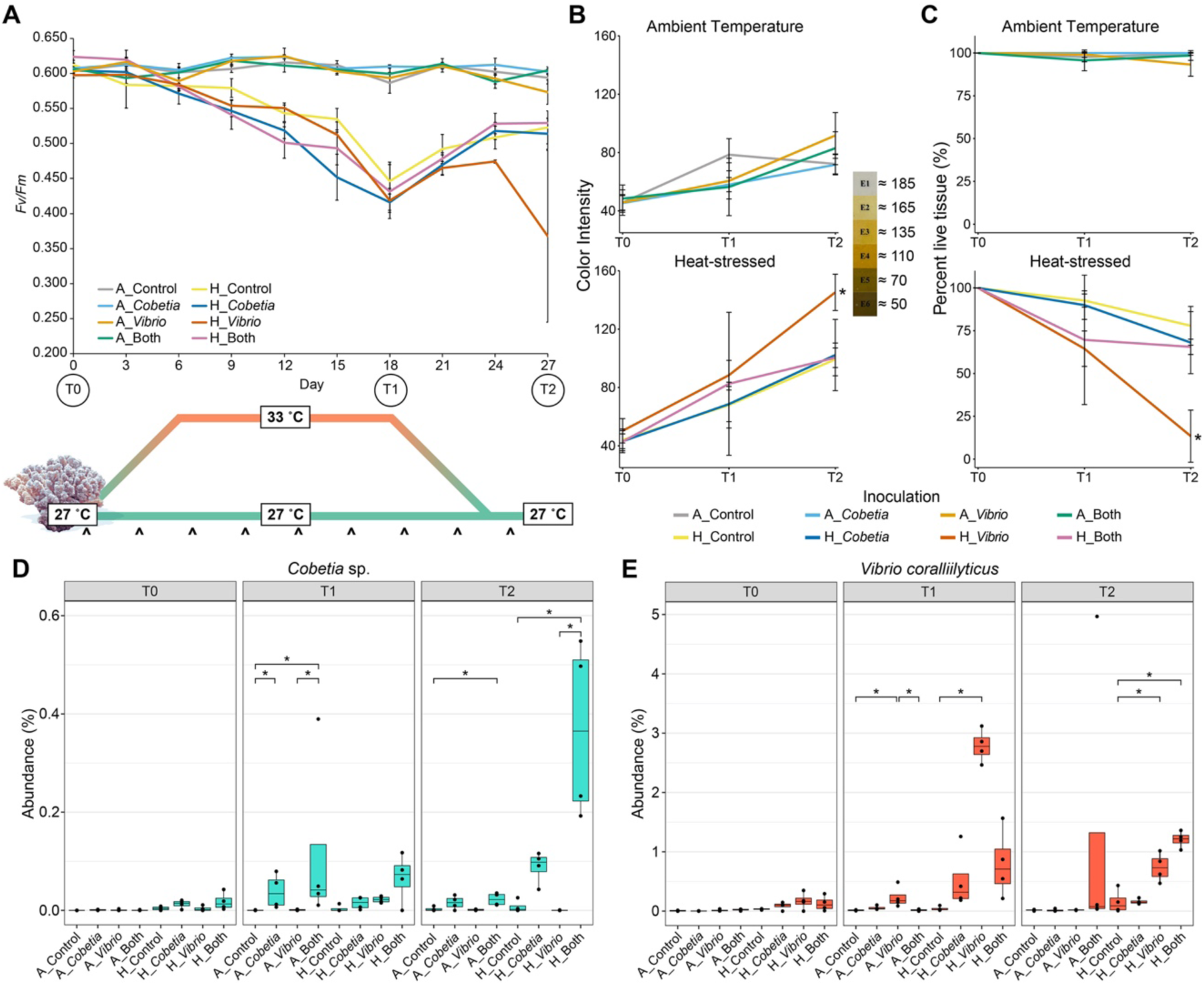
Coral phenotypic response and inoculated bacteria abundances throughout the experiment. (A) Mean photosynthetic efficiency (*Fv/Fm* ratios; y axis) from coral nubbins during the mesocosm experiment days (x axis). The error bars represent standard deviations (n = 4). Carets (^) below the schematic represent days of bacteria inoculation (every 3 days starting from day 1). (B) Average color intensity and (C) percentage live tissue of the coral nubbins in each condition, as measured by ImageJ (T0: n = 12 per group; T1/T2: n = 4 per group). Lines represent the mean value for each condition, and the error bars represent standard deviations. Higher color intensities represent more paling, bleaching, or tissue loss. Color intensity was measured on the CoralWatch Coral Health Chart^21^ for reference. Percentage abundances are shown in boxplots of the bacterial inocula (*Cobetia* sp. (D) and *V. coralliilyticus* (E)) across inoculations and sampling times (n = 4 per group). Boxplots display the individual data points, medians, lower and upper quartiles, and whiskers displaying the 1.5x interquartile range. Abundances of each inoculum were compared between each inoculation group (i.e., Control, *Cobetia*, *Vibrio*, Both) within each temperature condition (i.e., ambient temperature or heat-stressed). Asterisks denote significant differences between groups within a timepoint (*Dunn test*: p < 0.05).

Color intensity of coral tissue was measured by converting each image to grayscale and calculating the average intensity (from 0 = black, to 256 = white) across the whole image of the coral nubbin. Thus, a lower average color intensity would represent darker, i.e., healthier, living tissue, and a higher color intensity would represent pale, bleached, or dead coral tissue. Similar to previous findings^4^, the combination of heat stress and *Vibrio* inoculation (H_*Vibrio*) resulted in significantly higher color intensity, i.e., ‘whiteness’ or paling, (*Kruskal-Wallis*: *X^2^*= 8.49, df = 3, p = 0.037; *Dunn test*: p = 0.043; Fig. 1B) and lower percentage live tissue (*Kruskal-Wallis*: *X^2^* = 10.08, df = 3, p = 0.018; *Dunn test*: p = 0.014; Fig. 1C) when compared to the control group (H_Control) after the recovery period (T2) (Supplementary Table 1b). Notably, both the color intensity and percentage of live tissue in all other inoculation groups within the same temperature regime, including the co-inoculation group, were statistically indistinguishable from each other (Fig. 1B-C). Although the *Vibrio*-inoculated coral nubbins were paler than the nubbins from the other experimental groups at T2 in the heat-stressed corals, the color intensities of the live tissue remaining on the coral nubbins were not significantly different (*Kruskal-Wallis*: *X^2^* = 7.48, df = 3, p = 0.058; Extended Data Fig. 1).

Using 16S rRNA gene analysis, we found that the relative abundance of *V. coralliilyticus* was significantly higher in the corals treated with the pathogen than the control group at T1 (*Kruskal-Wallis*: *X^2^* = 13.059, df = 3, p = 0.005; *Dunn test*: p = 0.002) and T2 (*Kruskal-Wallis*: *X^2^*= 11.647, df = 3, p = 0.009; *Dunn test*: p = 0.045) (Fig. 1D-E and Supplementary Table 2). The abundance of *Vibrio* in the co-inoculation groups were lower than the *Vibrio*-inoculations at T1, but not at T2, although these differences were only significant between A_*Vibrio* and A_Both at T1 (*Kruskal-Wallis*: *X^2^* = 11.978, df = 3, p = 0.007; *Dunn test*: p = 0.023). This coincided with increased microbiome alpha diversity in all heat-stressed groups at T2 (Extended Data Fig. 2) and microbiome restructuring in the heat-stressed groups at T1 and T2 marked by a decrease in members of the family Endozoicomonadaceae, and increases in Saprospiraceae and Rhodobacteraceae (Extended Data Fig. 3). Significant differences in beta diversity were also observed between temperature groups and between the bacterial inoculation groups within both temperature conditions at T1 and T2 (Extended Data Fig. 4, Supplementary Table 3a and 3b).

### Coral epigenomes respond to temperature stress and bacterial inoculation

To explore whether the stress mitigation in co-inoculated coral nubbins was also attributed to interactions with the coral epigenome, genome-wide differences in DNA methylation were analyzed in response to temperature stress, bacterial inoculation, and their interaction. The profile of median methylation levels of genes across the genome of *P*. *verrucosa* were visualized with a non-metric multidimensional scaling (nMDS) plot for each timepoint (Fig. 2, Supplementary Table 4a). Significant differences existed at the peak of stress (T1) between corals exposed to the heat stress and those that were not (*PERMANOVA*: *R^2^* = 0.107, *F* = 3.245, p = 0.001; *PERMDISP*: p = 0.981), showing that methylation marks can change after twelve days of constant exposure to temperature stress (and 19 days of experimentation).

**Fig. 2.**
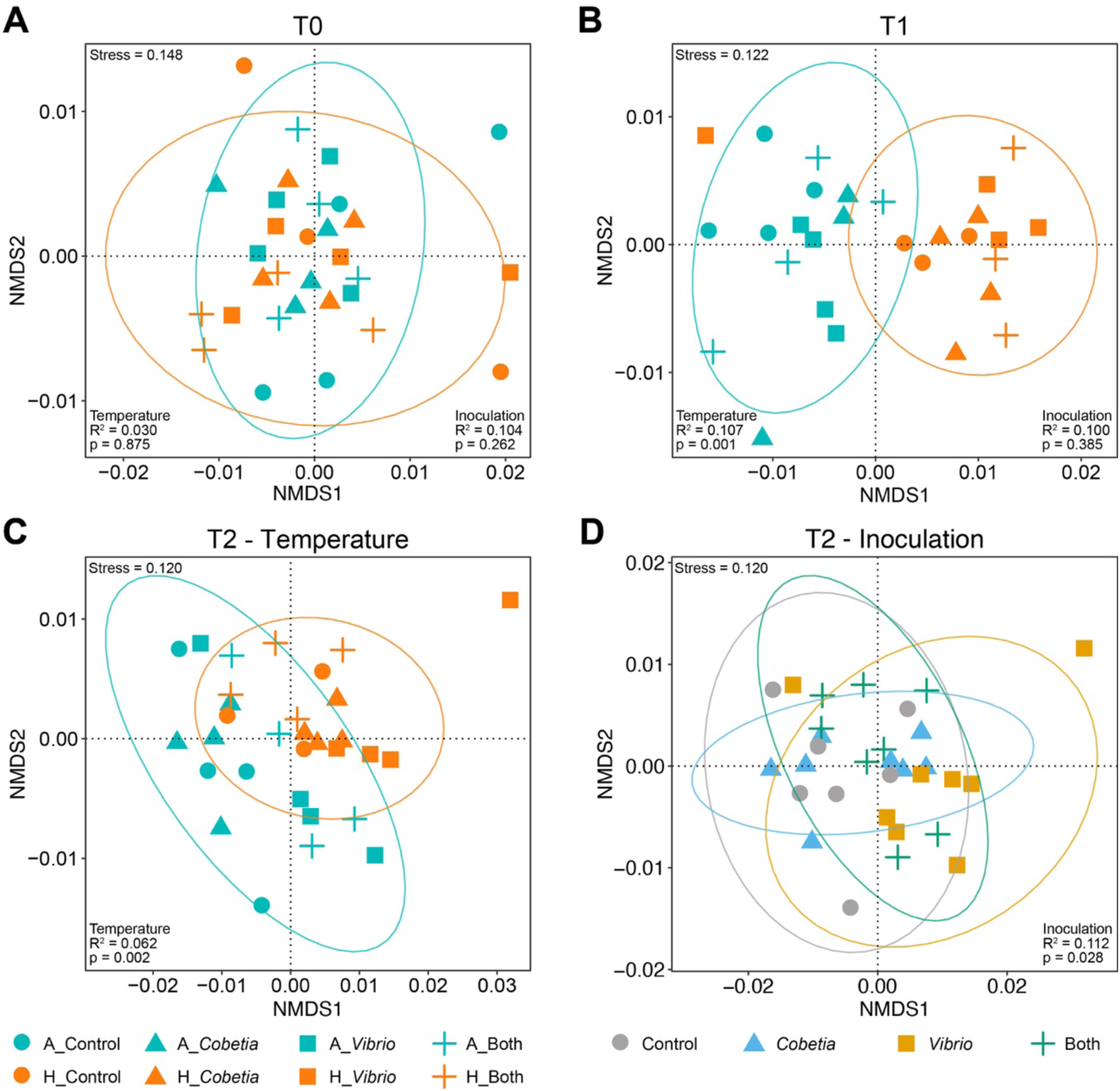
Non-metric MultiDimensional Scaling (nMDS) plots based on Bray-Curtis dissimilarities of the median methylation levels of genes across the genome of *P. verrucosa*. Samples from each timepoint were separated using Bray-Curtis dissimilarity and plotted along the first two dimensions. Circles = 95% confidence intervals around temperature groups. Both temperature and inoculation were tested as factors driving the separation between groups using PERMANOVA at (A) T0, (B) T1, and T2. R^2^ and p-values are displayed, representing the magnitude and significance attributed to each of the factors (n = 4 per group, except A_Control at T1, H_Control at T0,/T1/ T2, and H_Both at T1, where n = 3). nMDS plots for T2 are shown highlighting differences based on (C) temperature and (D) bacteria inoculation.

The difference between samples subjected to different temperature conditions was also observed after 9 days of recovery (28 days total; T2) (*PERMANOVA*: *R^2^* = 0.061, *F* = 1.955, p = 0.002; *PERMDISP*: p = 0.399). Additionally, ‘time’ was a significant factor in separating the heat-stressed corals using genome-wide DNA methylation levels, and pairwise comparisons showed significant differences between each of the timepoint pairs T0-T1, T1-T2, and T0-T2 (Extended Data Fig. 5). Bacterial inoculation groups also significantly contributed to the separation of samples at T2 (*PERMANOVA*: *R^2^* = 0.112, *F* = 1.197, p = 0.028; *PERMDISP*: p = 0.412), but there was not statistical support for the interaction between temperature and bacteria inoculations (*PERMANOVA*: *R^2^* = 0.106, *F* = 1.133, p = 0.106). Pairwise comparisons showed that pathogen-inoculated corals were significantly different from the control corals (*PERMANOVA*: *R^2^* = 0.101, *F* = 1.456, p = 0.029), but there was no statistical support for pairwise differences between any of the other inoculation groups at T2 (Supplementary Table 4b).

### Pathogen inoculation leads to changes in gene methylation

The significant genome-wide differences in median methylation levels observed between the temperature conditions at T1 and the temperature conditions and bacteria inoculation groups at T2 (Fig. 2) were further investigated using pairwise generalized linear models (GLMs). A total of 932 genes were significantly associated with temperature differences at T1, and 313 genes were significantly associated with the different temperature groups at T2 (Extended Data Fig. 6, Supplementary Table 5a and 5b). At T2, 17 genes were found to be significantly influenced by *Vibrio* inoculation in comparison to the control group, 15 of which had depleted median methylation levels correlated with *Vibrio* inoculation (Supplementary Table 5c). Extending the comparison of these 17 genes to the other inoculation groups using hierarchical clustering based on Euclidean distances shows that coral nubbins inoculated with *Vibrio* separated from the other three conditions at T2 (Fig. 3). These results corroborate the stressed physiological response of the coral nubbins caused by *Vibrio* inoculation observed at T2 (Fig. 1B-C) because it shows that the other inoculation groups (*Cobetia* and Both) cluster more closely to the Control group than the *Vibrio* inoculation in these genes. This demonstrates that, although pathogen inoculation significantly alters coral DNA methylation, co-inoculation of the pathogen and BMC leads to methylation patterns similar to the control group, supporting the notion that BMCs negate impacts caused by pathogen stress on coral health.

**Fig. 3.**
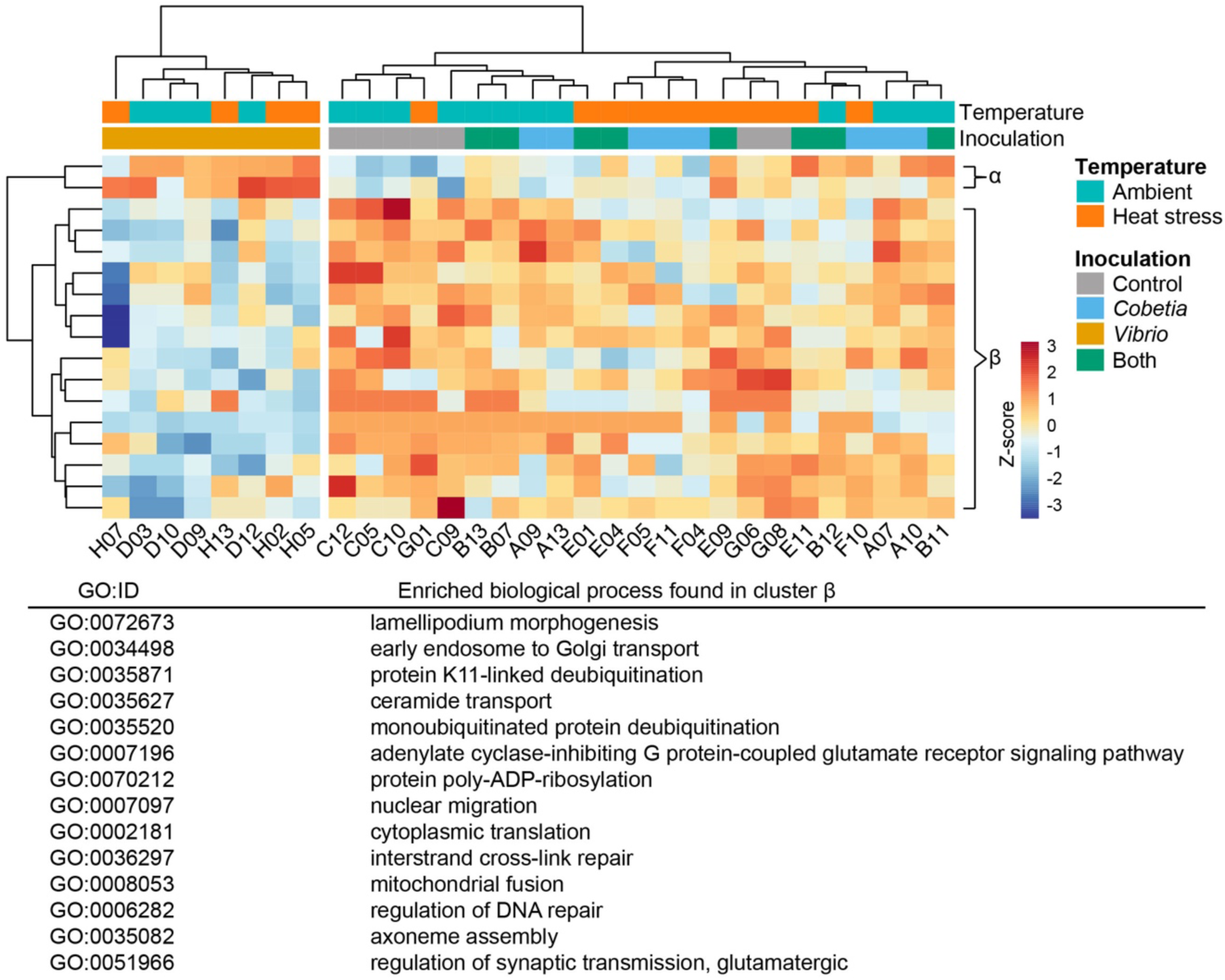
Genes with median methylation levels significantly associated with *Vibrio* inoculation according to GLMs compared to samples from other inoculation groups at T2 (n = 4; n = 3 for H_Control). There is a separation between the corals inoculated with *V. coralliilyticus* and the other inoculation groups (x-axis). Hierarchical clustering was performed using Euclidean distances and the “ward.D2” agglomeration method. The heights of the dendrograms, representing distances separating the clusters, ranged from 2.6 to 18.3 between the coral samples and 4.0 and 14.1 between the methylated genes. Colors represent the z-score for any given gene. The enriched GO terms using TopGO display the significant functions with depleted methylation levels in pathogen-stressed corals (cluster β). All enriched GO terms are in Supplementary Table 6c. All GO terms associated with these genes are in Supplementary Table 7.

Increases in gene body methylation reduce spurious transcription in highly expressed genes, thereby reinforcing the epigenetic control of genes as expression increases^14–16^. Our findings here revealed that methylation within *P*. *verrucosa* occurred primarily within the gene body (Extended Data Fig. 7, Supplementary Table 8). Genes with depleted methylation in the corals treated with *Vibrio* only (cluster β in Fig. 3) highlight the link between pathogen stress and diminished epigenetic control of genes involved in cell maintenance and functioning and genes related to the immune system, which potentially correlates with decreased gene expression.

The enriched GO terms in cluster β included “adenylate cyclase-inhibiting G protein-coupled glutamate receptor signaling pathway” and the “regulation of synaptic transmission (glutamatergic)”, which both rely on the neurotransmitter glutamate.

Glutamate production is necessary for algal symbiosis in the cnidarian model *Aiptasia*^22^. Moreover, in the coral *Stylophora pistillata*, heat stress was found to disrupt glutamate biosynthesis and trigger glutamate catabolism, which contributed to coral bleaching^23^.

Therefore, pathogen inoculation may exacerbate losses in glutamate caused by heat stress by reducing the epigenetic control of glutamatergic signal transduction. This may be further supported by significant differences at T2 in the GO term “glutamatergic synapse” (GO:0098978) due to temperature conditions (Supplementary Table 6b), although additional research in this area is necessary, as heat stress did not induce uniform methylation trends in all of the affected glutamatergic synapse genes (Extended Data Fig. 8). Further coral immune functions with depleted median methylation levels due to *Vibrio* inoculation include “lamellipodium morphogenesis”, “early endosome to Golgi transport”, and the stress response mechanisms “interstrand cross-link repair” and “regulation of DNA repair”.

### Pathogen inoculation affects gene expression

To determine whether the methylation changes correlated with changes in gene expression levels, the transcriptomes of the T2 nubbins were analyzed using RNA-Seq. Heat stress and pathogen inoculation caused distinct transcriptome changes, with H_*Vibrio* samples separating from the other groups in a principal component analysis (PCA) plot along PC1 (which accounted for 74.93% of the variance) (Fig. 4A).

**Fig 4.**
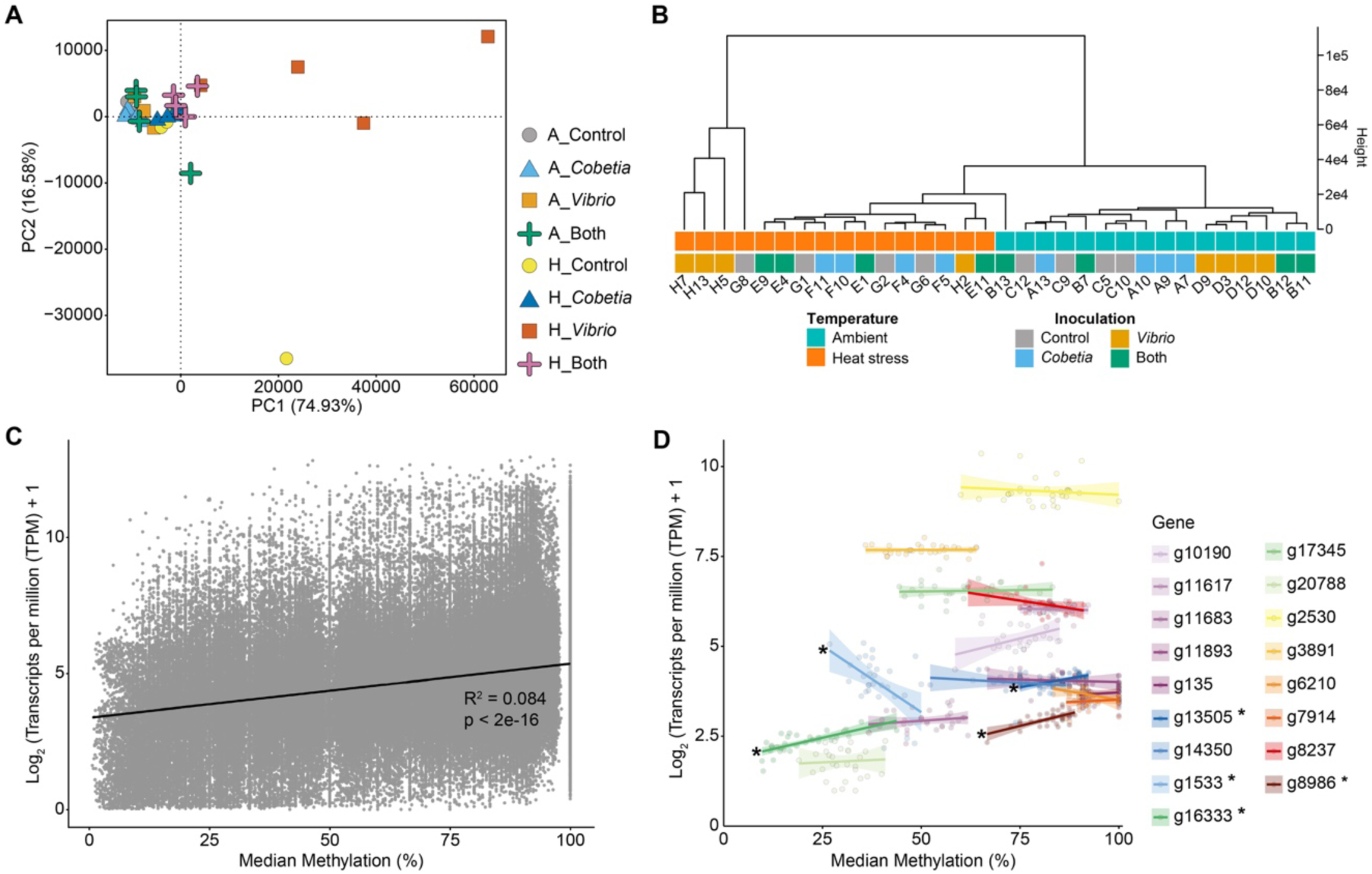
Gene expression levels are affected by heat stress and bacteria inoculation and correlated with methylation levels at T2. (A) PCA generated from the transcripts per million (TPM) expression estimates of *P*. *verrucosa* genes (n = 4). (B) Hierarchical clustering using the “ward.D2” agglomeration method to separate TPM values across the samples (n = 4). (C) A linear model shows that median methylation levels for all of the genes across all 31 samples were significantly and positively correlated with gene expression (using Log_2_ (TPM + 1). The R^2^ and p-value here represents the magnitude and significance attributed to the correlation. (D) The 17 genes with median methylation percent affected by pathogen inoculation are plotted against their respective gene expression. Four of these genes had median methylation levels, which significantly correlated with gene expression (denoted with asterisks).

Hierarchical clustering also showed that three of the four H_*Vibrio* samples separated distinctly from the other groups (Fig. 4B). The median methylation levels across the genome positively correlated with the relative expression level of genes (R^2^ = 0.084, p < 2e-16; Fig. 4C). Of the 17 genes with altered methylation due to pathogen inoculation, four showed a significant correlation between changes in methylation and gene expression across experimental groups (Fig 4D and Supplementary Table 9). Three of the genes: g13505, g16333, and g8986, were positively correlated with gene methylation, and were defined by the GO terms related to transmembrane transporter activity, negative regulation of transcription by RNA polymerase II, and cilium movement, respectively (Supplementary Table 7). One gene, g1533, was negatively correlated with gene methylation and was not associated with a GO term (Supplementary Table 7).

### Gene methylation differences are correlated with phenotypic response

To determine which genes affected by *Vibrio* inoculation (Fig. 3) correlated most with the paling and tissue loss observed in the corals (Fig. 1), linear models were implemented using the median methylation levels and color intensity as factors (Supplementary Table 10). The genes showing the highest correlation between methylation percentage and phenotypic response (R^2^ > 0.100 and p < 0.05) were g16333, g1533, g17345, g14350, and g8986 (Fig. 5). Example GO terms describing these genes were “negative regulation of transcription by RNA polymerase II” (GO:0000122), N/A, “ceramide transport” (GO:0035627), “adenylate cyclase-inhibiting G protein-coupled glutamate receptor signaling pathway” (GO:0007196), and “cilium movement” (GO:0003341), respectively (Supplementary Table 7). Of the genes with defined functions, three are related to cellular processes, while g17345 (“ceramide transport”) functions in macromolecule localization within the cell. In all of the defined genes, *Vibrio*-inoculation led to a significant depletion in DNA methylation when compared to the control group in either the ambient temperature or heat-stressed conditions. In all but one gene (g16333), co-inoculation with *Cobetia* prevented similar decreases in DNA methylation when compared with the control group. The genes g14350, defined by the GO term “adenylate cyclase-inhibiting G protein-coupled glutamate receptor signaling pathway”, and g1533, had significantly different median methylation levels between *Vibrio*-treated corals and the co-inoculation in the heat-stressed condition (*ANOVA*: *F* = 4.46, df = 3, p = 0.028, *TukeyHSD*: p = 0.033 for g14350; *ANOVA*: *F* = 11.62, df = 3, p = 0.001, *TukeyHSD*: p = 0.014 for g1533).

**Fig. 5.**
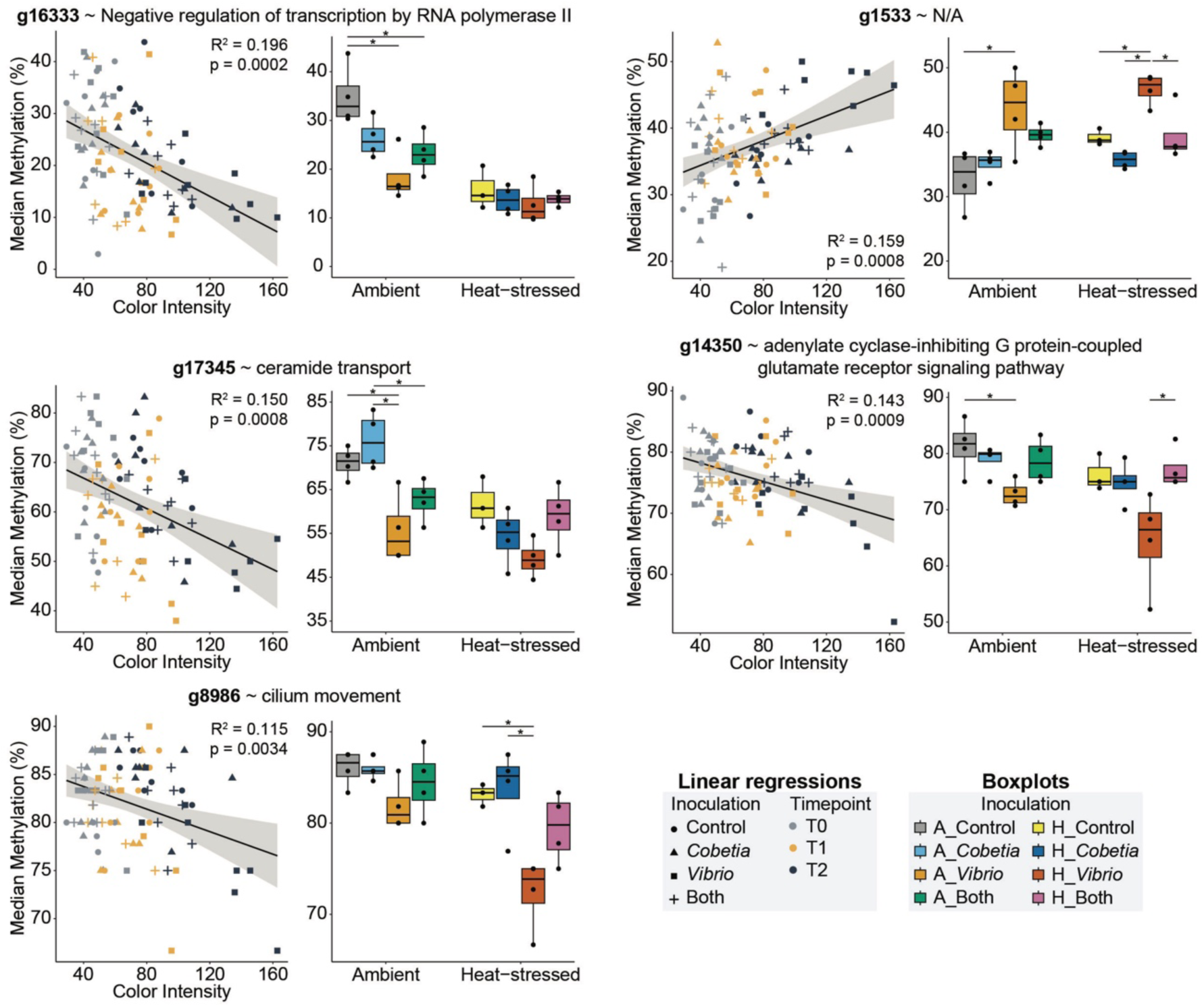
Methylation profiles of genes with significant correlations (R^2^ > 0.100, p < 0.05) between methylation levels and phenotypic response (coral nubbin color intensity). Linear regression plots show the correlation lines of the coral nubbin color intensity to the median methylation percent in each sample (n = 91) along with the 95% confidence intervals (shaded regions). Boxplots showing the median methylation percent of the samples at T2 (after recovery), were used to compare between groups with one-way ANOVA, followed by TukeyHSD (n = 4; n = 3 for H_Control). Asterisks denote significance between inoculation groups in the boxplots (p < 0.05). Boxplots display the individual data points, medians, lower and upper quartiles, and whiskers displaying the 1.5x interquartile range.

## Discussion

Although hypothesized, the interaction between changes in the coral-associated microbiome and the coral epigenome was yet to be demonstrated^13,18^. The findings of this study are fourfold, as we show that, in corals, (1) the addition of a single probiotic (*Cobetia* sp.) was able to reduce the indicators of stress caused by inoculation with the pathogen *V*. *coralliilyticus*; (2) bacterial inoculation, as well as temperature stress, contribute to differences in DNA methylation; (3) gene expression is, on the whole, positively correlated with methylation levels and affected by pathogen and heat stress; (4) pathogen inoculation causes changes in DNA methylation and gene expression to specific cellular processes and immune responses that was mitigated by the probiotic.

Regarding the phenotypic response, differences in color intensity and tissue loss observed in the coral nubbins at T2 between the control group and the *Vibrio* inoculation in the heat-stressed corals may be a delayed effect of the establishment of the pathogen in the heat-stressed samples at T1. Interestingly, the relative abundances of *V*. *coralliilyticus* was lower at T2 than T1, coinciding with the observed tissue loss. This result, and the changes to bacterial community diversity indices in heat-stressed nubbins indicate that the holobiont is becoming increasingly dysbiotic^24^. Furthermore, the co-inoculation with the BMC mitigated *Vibrio* establishment in the microbiome during the peak of stress at T1, and controlled the relative abundances of *Vibrio* to similar degree at T2. The co-inoculation also had a positive impact on the phenotype in heat-stressed corals at T2, contributing to the increased stress resilience and survival observed in this group versus the *Vibrio*-inoculated group. Together, this supports the notion that, although heat stress leads to overall changes in the microbial community, attenuation of the pathogen by the addition of probiotics boosts coral resilience.

DNA methylation has been shown to be responsive to environmental changes^15,17,25–28^, correlate with phenotypic acclimatization^15,28^, and be heritable^29^. Plasticity of DNA methylation in corals thus far has been tested on the scale of weeks^28,30–32^ and months^17,26,27,33^ to a year or more^14,15,25,33^. The findings here further prove the plasticity of coral epigenomes during stress, as 12 days of sustained heat stress induced DNA methylation changes across the genome. Furthermore, the difference continued during a nine-day recovery period, which exemplifies persistence of epigenome modifications brought about by environmental change.

Recently, the concept of “epigenetic memory”, where intragenerational differences between epigenomes of different individuals lead to vertical transmission of distinct epigenomes, was confirmed in the brain coral *Platygyra daedalea*^29^. This concept has been recognized in plants for many years (reviewed in ^34^). Here, the changes in epigenomes driven by environmental differences as well as bacterial inoculations, signify the potential for long-term persistence after removal of the environmental stress and shows promise for microbiome manipulation experiments with implications for long-term acclimation and coral resilience.

When analyzing the methylation levels of the genes and putative functions, we can conclude that *Vibrio* inoculation caused changes to methylation levels in metabolic and cellular processes relevant to the observed phenotypic responses (Fig. 1). The difference between the *Vibrio*-inoculated corals and the other three inoculation groups (control, inoculation with *Cobetia* sp., and the co-inoculation) in these genes at T2, independent of the temperature regime, demonstrates the coral probiotic’s ability to prevent changes to the coral epigenome in key functions altered by the pathogen (*V*. *coralliilyticus*). This effect was associated with increased resilience to stress and higher survival rates.

Moreover, methylation of functions related to algal symbiosis (e.g., “adenylate cyclase-inhibiting G protein-coupled glutamate receptor signaling pathway” and “lamellipodium morphogenesis”) and immune responses (e.g., “early endosome to Golgi transport”) were depleted in *Vibrio*-inoculated corals. As mentioned above, pathogen stress may reduce epigenetic control over glutamate receptor pathways, disrupting algal symbiosis and compromising coral health. Glutamate mediates interactions between corals and their algal symbionts^22^, and inhibiting glutamate receptors disrupts this relationship^35^.

Lamellipodia, and by association “lamellipodium morphogenesis”, also contributes to coral-algal interactions^36^, as well as wound healing in the cnidarian *Clytia hemisphaerica*^37^. Reduced methylation of “early endosome to Golgi transport” supports previous findings that *Vibrio* targets endosomal pathways during infection^38^. Specifically, toxins produced by *V*. *coralliilyticus* disrupt endosome development and downregulate immune genes in the host^38,39^. These methylation changes indicate the *Vibrio*’s ability to elicit a response to coral host defense genes, and the lack of difference between co-inoculated corals and the other inoculations shows the probiotic’s ability to mitigate this effect.

We also confirmed that overall gene expression was positively correlated with gene methylation, which adds to the growing evidence that basal coral epigenomes and gene expression levels are correlated^14,15,31,40^. The correlated down-methylation and down-regulation of genes related to transmembrane transporter activity (g13505), cilium movement (g8986), and negative regulation of transcription by RNA polymerase II (g16333) due to pathogen exposure may indicate direct mechanisms or host responses to the stress. For example, transmembrane transporters are essential for transporting nutrients between corals and their symbionts; reduced expression of related genes may signal a breakdown in symbiosis^41^. The pathogen’s impact on cilium movement expression and methylation also shows that the stressed coral nubbins may have been unable to remove excess reactive oxygen species^42^, which could have contributed to tissue bleaching and death observed in our samples. RNA polymerase II, on the other hand, helps control spurious transcription by recruiting the histone-methylating enzyme SetD2 to transcribed DNA^14,16^. Therefore, pathogen-induced repression of this gene demonstrates a disruption in both the transcriptional and epigenetic control of gene fidelity. The significant positive correlations between DNA methylation in the latter two genes (g8986 and g16333), as well as in the glutamate receptor pathway gene (g14350), and the coral’s phenotypic response to *Vibrio* inoculation further demonstrates the importance of integrating coral epigenomes into coral immune research.

Despite the association between DNA methylation and gene expression, questions remain about the direction and magnitude of this relationship, and its implications for acclimation and adaptation ^31,32^. This has led to the hypothesis that coral gene methylation responds to transcription levels, rather than functioning as a gene regulatory mechanism like it does in mammals ^14,27,32^. Thus, epigenomes may help to explain how rapid gene expression changes caused by pathogen stress lead to long-term phenotypic responses.

The direct mechanisms by which bacteria are able to elicit changes to the coral epigenome are still to be explored, although metabolites and proteins produced by coral-associated bacteria are suggested to play a role^13,18^. Additionally, the crosstalk between coral epigenomes and coral-associated bacteria may not be unidirectional, as pathogen susceptibility has been hypothesized to be at least partly controlled by epigenetic factors in aquaculture fish^43^. By connecting rapid microbiome changes to more long-term and heritable epigenetic changes (Fig. 6), bacteria inoculation has the potential to induce lasting differences, even without direct establishment in the bacterial community. Thus, as future studies explore this interaction further, the current paradigm that microbiome interventions only lead to short-term and volatile changes^44^ might shift. Therefore, as research in this field progresses, there is considerable potential to shape the use of customized microbial therapies for increasing resistance and resilience in vulnerable corals^45,46^.

**Fig. 6.**
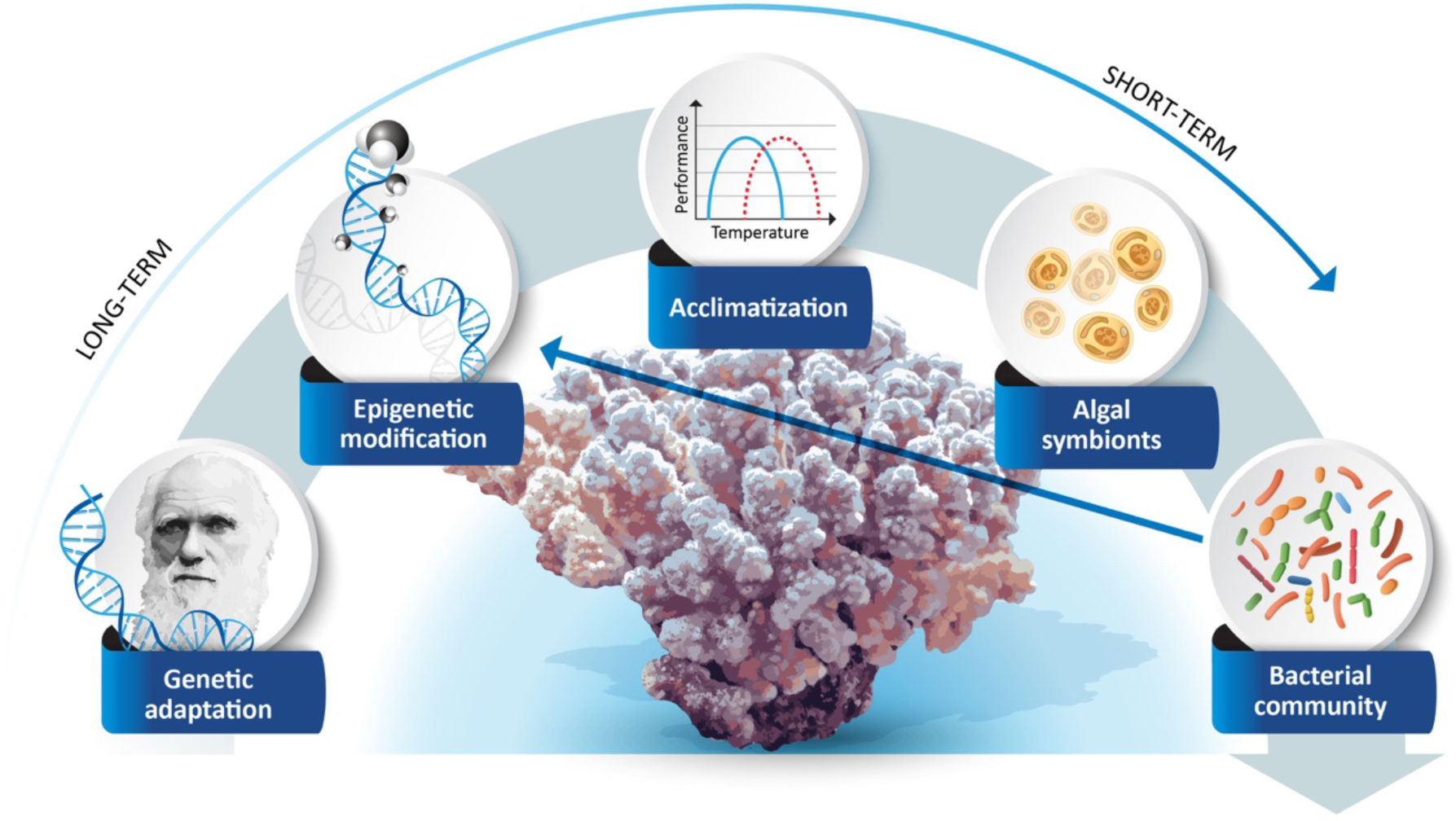
Rethinking the flexibility and longevity of coral microbiomes via crosstalk with coral epigenomes. Microbiome changes occur quickly and are fleeting. However, relating microbiome manipulation and coral DNA methylation here shows that bacterial community changes may induce long-term and possibly heritable changes in corals by influencing modifications to the coral epigenome. This figure highlights the transformative nature of our results by adding an arrow to the previous concept postulated in ref.^47^, connecting the bacterial community with epigenetic modification.

## Methods

### Coral collection and experimental setup

In order to prevent intergenomic variability would obfuscate identification of methylation marks^48^, clonal fragments from *Pocillopora verrucosa* were collected from the Al Fahal reef in the central Red Sea (22° 18.3071’ N, 38° 57.8741’ E) on 14 March 2022 (*in situ* seawater temperature = 25°C) and brought to the aquariums at the Coastal and Marine Resources Core Lab (CMR) in King Abdullah University of Science and Technology (KAUST). All coral husbandry and experimentation was done under the supervision of KAUST’s Institutional Biosafety and Bioethics Committee (IBEC). The coral colony was acclimated to CMR conditions (using a 300 L open-system aquarium tank, with a maximum intensity of 180 μmol of photons m−2 s−1 between 10:00 to 14:00, and a 12-hour day/night cycle changing at 7:00 and 19:00) and brought to 27°C over the course of three weeks (approximately 0.5°C increase every two days, followed by two weeks at 27°C). Corals were further fragmented into finger-sized nubbins on 4 April 2022. After a five-week recovery period, nubbins were divided into the eight experimental tanks which housed the replicates for each inoculation group (10 L aquarium tanks, using the same light parameters as the acclimation tank above) containing 0.22 µm filtered seawater. The coral nubbins were acclimated for seven days (until the average *Fv/Fm* ratio, a measure for photosynthetic efficiency and proxy for coral health, from the coral nubbins in each experimental tank was >0.600, using a Diving-PAM II; Walz, Germany).

The bacteria used in the experiment were the BMC *Cobetia* sp. strain pBMC4 (isolated from *Pocillopora verrucosa*^19^) and the opportunistic pathogen *Vibrio coralliilyticus* strain BAA-450 (isolated from *Pocillopora damicornis*^20^). The bacteria cultures used in this experiment were grown using Marine Broth 2216 (Becton Dickinson, USA) for 24 hours at 27°C before each inoculation. To minimize additional carbon input from the culture media, the cultures were spun at 6000 x g for 5 min and the bacteria pellets were resuspended in sterilized 3.5% NaCl solution. This process was repeated for a total of two washes, then the bacteria were resuspended in sterilized 3.5% NaCl solution for inoculation. Abundance and purity of each of the cultures were checked by the plate counting method using Marine Agar 2216 (Becton Dickinson, USA).

Starting on day 1, bacteria were inoculated into each of the tanks every three days at a final concentration of 10^5^ cells/mL. Thirty percent of the water was replaced with new 0.22 µm filtered seawater (kept at the same temperature of the experimental tanks) every other day. The experiment was performed over 28 days. Of the eight experimental tanks, four were maintained at 27°C throughout the entirety of the experiment (denoted with an “A” for ambient temperature). In the other four experimental tanks, starting from day 1, temperature was increased from 27°C to 33°C (by increasing the temperature 1°C per day for 6 days), then maintained at 33°C for 12 days (until the average *Fv/Fm* across all conditions was <0.450), then returned to 27°C (by decreasing the temperature 1°C per day) where they were kept until the end of the experiment (4 days). These tanks are denoted with an “H” for heat stress. The four experimental tanks in each of A or H, contained one of the following experimental conditions: corals with no inoculation (“Control”), corals inoculated with *Cobetia* sp. at a final concentration of 10^5^ cells/mL (“*Cobetia*”), corals inoculated with *V. coralliilyticus* at a final concentration of 10^5^ cells/mL (“*Vibrio*”), and corals inoculated with both *Cobetia* sp. and *V. coralliilyticus*, each at a final concentration of 10^5^ cells/mL (“Both”). In total the eight conditions were: A_Control, A_*Cobetia*, A_*Vibrio*, A_Both, H_Control, H_*Cobetia*, H_*Vibrio*, and H_Both.

Samples of four coral nubbin replicates (n = 4) per tank were taken at the start of the experiment before any bacterial inoculation (T0), on day 19 (peak of stress, T1), and on day 28 (after recovery, T2). At each sampling time, the apical tip of the sampled nubbin was removed and flash frozen in liquid nitrogen for nucleic acid analysis. Frozen nubbins were then transferred to -80°C until processing.

### Coral health parameters

Coral health was monitored during the experiment using a Diving-PAM II (Walz, Germany) to measure the *Fv/Fm* ratios. *Fv/Fm* measurements were taken after dark acclimating the coral nubbins for at least 1 hour the night before each inoculation day (every three days starting on day 0) throughout the entirety of the experiment (n = 4 for each experimental tank). *Fv/Fm* measurements were compared between temperature groups using Wilcoxon Rank Sum tests^49^, and between inoculation groups Kruskal-Wallis H tests^50^. Images were taken of the coral nubbins at T0 and at their respective sampling times (T1 or T2).

ImageJ was used to analyze the percent live tissue, color intensity, and color intensity of live tissue of each of the images of the nubbins from sampling times T0 (n = 12 for each group), T1 (n = 4 for each group), and T2 (n = 4 for each group). For the percent live tissue, the area of live tissue on the coral nubbin was divided by the total area of the coral nubbin in each respective image. The color intensity, as described in the main text, was obtained using the unweighted average in color intensity for each coral nubbin, after converting the image to 8-bit grayscale (offering 2^8^ = 256 levels of differentiation). The average color intensity incorporates the color intensities (from 0 = black, to 256 = white) of all pixels within a coral nubbin. This, then, incorporates the color of the live tissue and any exposed coral skeleton that may be present in coral nubbins with tissue loss. For the color intensity of live tissue, only the unweighted average in color intensity observed in the live tissue portion of the image was considered. Percent live tissue, color intensity, and color intensity of live tissue measurements were compared between the samples with Kruskal-Wallis H tests^50^, followed by Dunn’s multiple comparison test^51^ using Benjamini-Hochberg adjustment.

### 16S rRNA gene amplicon sequencing

DNA and RNA were extracted from airbrushed coral tissue using AllPrep DNA/RNA Mini Kit (Qiagen, Germany), following manufacturer’s instructions. Extracted DNA was used for analysis of the bacterial community as well as identification of the methylation marks within the coral genome. For bacterial community analysis, the DNA composing the V3-V4 region of the 16S rRNA gene was sent to Novogene (Novogene, China) for sequencing using a NovaSeq 6000 (Illumina, USA) with the primers 341F (5’-CCTAYGGGRBGCASCAG-3’) and 806R (5’-GGACTACNNGGGTATCTAAT-3’).

### Library preparation and sequencing of methylated DNA

Libraries of converted methylated DNA were prepared using the NEB Enzymatic Methyl-seq Kit (New England Biolabs Inc., USA) with the EM-seq Index Primers (New England Biolabs Inc., USA). Prepared libraries were sent to Novogene (Novogene, China) for sequencing using a NovaSeq 6000 (Illumina, USA). Due to insufficient DNA quality in 5 samples, 91 of the 96 samples were used for downstream analysis (n = 4 for each experimental group; except A_Control at T1; H_Control at T0, T1, and T2; and H_Both at T1; which all had n = 3).

### Coral microbiome analysis

For microbiome analysis, primer sequences were removed using cutadapt^52^. Sequences were further filtered and trimmed using the DADA2 pipeline^53^. Taxonomy was assigned using the SILVA_nr99_v138.1 database^54^. Sequences identified as Bacteria that did not match to Chloroplast or Mitochondria were further analyzed using phyloseq^55^. ASVs with a 100% match to the 16S rDNA sequence of the inoculums (*Cobetia* sp. or *Vibrio coralliilyticus*) via BLASTN^56^ were determined to be representative ASVs of the inoculated bacteria. Statistical differences between relative abundance data in different groups were determined using the *vegan* package in R (version 4.3.1). Bacterial abundances were not normally distributed (*Shapiro*-*Wilk*: p < 0.05), so significant differences between groups were determined by non-parametric tests. Kruskal Wallis H tests^50^, followed by Dunn’s multiple comparison tests^51^, were used to find significant differences between inoculation groups (n = 4 for each experimental group).

Microbiome composition differences were calculated between inoculation groups in each of the temperature conditions and timepoints using principal coordinate analyses (PCoA). Significant factors driving the separation between samples and sample dispersion were determined using the adonis2 and betadisper commands in the *vegan* R package (version 4.3.1). Pairwise differences between inoculation groups were tested using the pairwise.adonis command in the *pairwiseAdonis* R package.

### Coral DNA methylation

DNA methylation was analyzed bioinformatically following the steps described in Liew et al. (2020)^29^. Reads were first trimmed using TrimGalore v0.6.10^52^. Then, the filtered reads were mapped to a reference genome for *Pocillopora verrucosa*^57^ and deduplicated, before methylation marks were scored per-position using Bismark v0.23.1^58^. Methylated DNA positions were only considered bona fide if they met the criteria defined in Liew et al. (2020)^29^. Briefly, the probability of methylated positions resulting from sequencing error needed to be below 0.05 after correcting for false discovery rate, and the methylated reads were only kept if the methylation occurred in all replicates of at least one of the experimental conditions. Also, to minimize type I errors, the median coverage needed to be ≥ 10 and minimum coverage needed to be ≥ 5 across all of the samples in order to be considered for further analyses^14,15,29^ (Supplementary Table 11).

The median methylation levels across genes were compared between groups. Each methylated position within a given gene was given a score between 0 (no methylation in any of the cells) to 1 (fully methylated across all cells), and the median across each gene was used for further analyses. Only genes with five or more methylated positions were used for downstream analyses. Nonmetric multidimensional scaling (nMDS) plots were used to compare the median methylation across the genome of the coral using Bray-Curtis dissimilarity to separate samples. Significant factors driving the separation between samples and sample dispersion were determined by PERMANOVA using the adonis2 and PERMDISP using betadisper commands in the *vegan* R package (version 4.3.1). Pairwise differences between groups were tested using the pairwise.adonis command in the *pairwiseAdonis* R package.

To extrapolate the genes related to the difference observed between the Control inoculation and the *Vibrio*-treated corals at T2, generalized linear models (GLMs) were implemented in R using the following formula:

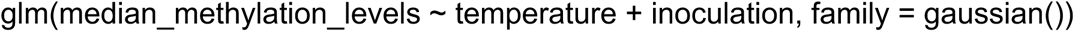

for the data from T2, where median methylation levels for each gene was compared with the predictor variables temperature (ambient vs. heat-stressed) and inoculation (by using the Control inoculation as a baseline to compare with each of *Cobetia*-treated, *Vibrio*-treated, and the co-inoculation inoculation samples). Genes with fewer than 5 methylated positions were removed from the analysis. Genes with *Vibrio* inoculation as a significant predictor variable (after correcting for false discovery rate) were plotted on a heatmap with all of the T2 samples using hierarchical clustering with Euclidean distances and the “ward.D2” agglomeration method.

### Functional enrichment of differently methylated genes

GO term annotations were collected from the published *P*. *verrucosa* genome^57^. GO term enrichment analysis was performed using TopGO^59^ on the genes with significantly different methylation levels from the previous step. Enriched GO terms with p < 0.05 and were considered significant.

### RNA-Seq analysis

T2 RNA samples (n = 32) extracted using the AllPrep DNA/RNA Mini Kit (Qiagen, Germany) were lyophilized onto a GenTegraRNA plate (Gentegra, USA) and sent to the MR DNA (Molecular Research LP, USA) for RNA-seq analysis. There, the lyophilized RNA was resuspended in 30uL RNase free water, and 500-750 ng of total RNA was used for library preparation using the poly-A selection method of KAPA mRNA HyperPrep Kits (Roche, Switzerland) following the manufacturer’s instructions. The libraries were then pooled in equimolar ratios of 0.6 nM and sequenced using a NovaSeq 6000 (Illumina, USA). RNA-seq reads were first trimmed using TrimGalore v0.6.10^52^. The adapter-trimmed reads were pseudo-aligned to *P*. *verrucosa* gene models^57^ using kallisto v0.50.1^60^ to quantify gene expression levels. Transcripts per million (TPM) obtained from sleuth v0.30.1^61^ were then used to compare across inoculation and temperature groups from T2 (n = 4 for each condition). Correlations between TPM and median methylation levels across genes were determined using the linear model:

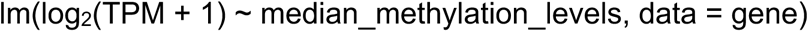

Significant correlations in individual genes were determined using Benjamini-Hochberg adjusted p-values. Gene expression levels across the T2 samples were then compared pairwise between the temperature groups (n = 16 per temperature regime) and inoculation groups (n = 8 per inoculation group).

### Correlating median methylation with phenotypic parameters

The genes predicted to be associated with *Vibrio* inoculation in the previous step were then tested against a phenotypic response variable using linear models with the formula:

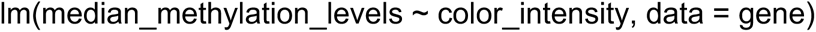

to determine which of these genes correlated with phenotypic differences observed in the corals affected by temperature and inoculation. The genes with R^2^ > 0.100 and p < 0.05 (after Benjamini-Hochberg correction) were plotted using linear regressions with 95% confidence intervals. Boxplots showing the median methylation levels of these genes in each sample at T2 were then used to compare differences between groups using one-way ANOVA followed by TukeyHSD.

## Data availability

EM-seq data is deposited in the NCBI Sequence Read Archive under the BioProject accession number PRJNA1123499 (https://www.ncbi.nlm.nih.gov/sra/PRJNA1123499).

## Code availability

The code used for data analysis can be found at: https://github.com/arbarno/pver_epibac.

## Acknowledgements

We would like to thank the Coastal and Marine Resources Lab and Bioscience Core Lab for their support and assistance throughout the experiment. This work was supported by KAUST grant number BAS/1/1095-01-01

## Author contributions

A.R.B., H.D.M.V., G.C., T.T., M.A., C.R.V., and R.S.P. designed the experiments; A.R.B., H.D.M.V., P.M.C. conducted the mesocosm experiment and sample collection; A.R.B. and N.D. performed DNA extractions; A.R.B. and F.C.G. performed analyses; A.R.B. wrote the manuscript with input from all authors; A.S.R. and R.S.P. provided guidance and funding during the experiment.

## Competing interests

The authors declare no conflicting interests.

## Materials & Correspondence

Correspondence to Adam R. Barno or Raquel S. Peixoto.

## Supplementary Information

Supplementary tables are provided in excel format.

**Extended Data Fig. 1.**
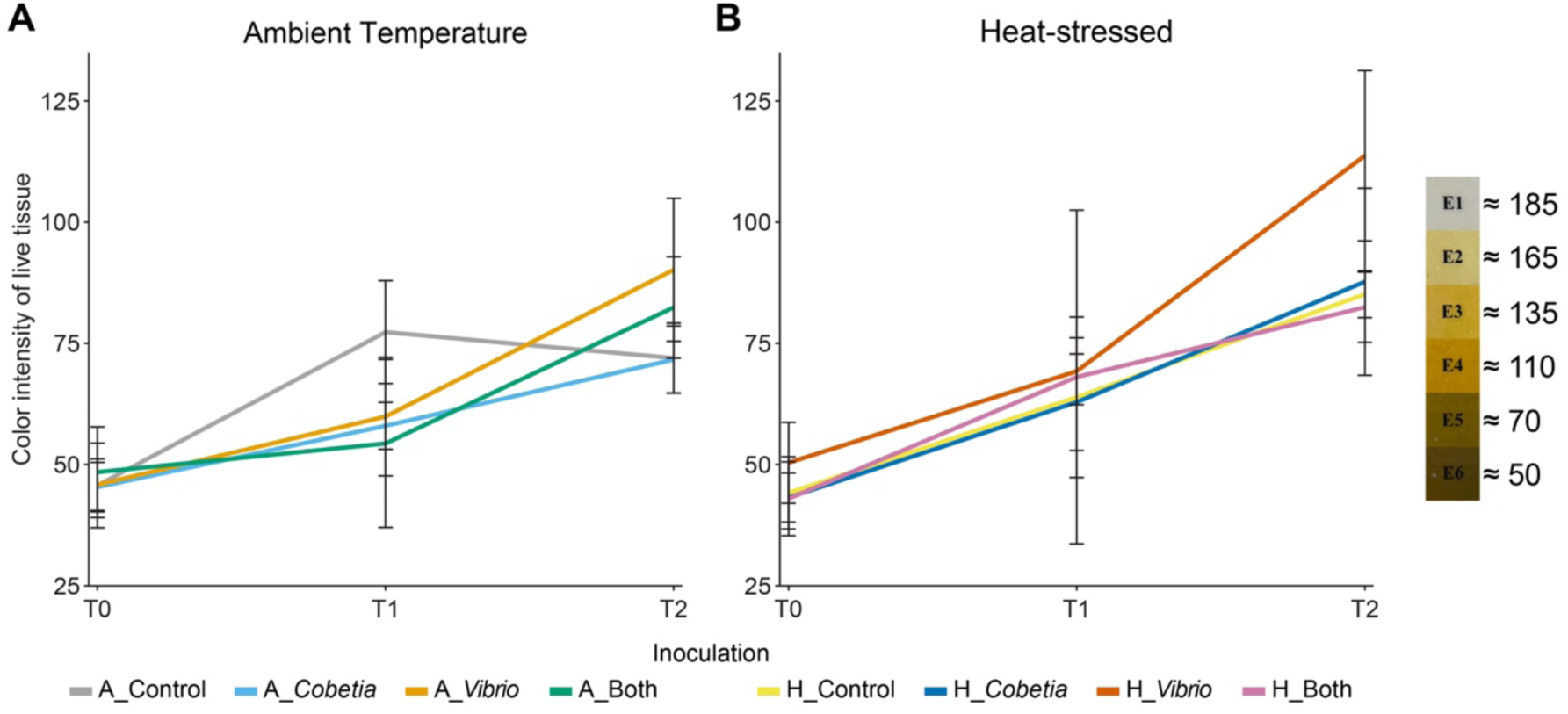
Color intensity of live tissue for the coral nubbins throughout the experiment. Lines represent the mean color intensities are plotted for each bacterial inoculation group at each timepoint for nubbins in the (A) ambient temperature condition and (B) heat-stressed condition, as measured by ImageJ (T0: n = 12 per group; T1/T2: n = 4 per group). The error bars represent standard deviations. Color intensity was measured on the CoralWatch Coral Health Chart^21^ for reference.

**Extended Data Fig. 2.**
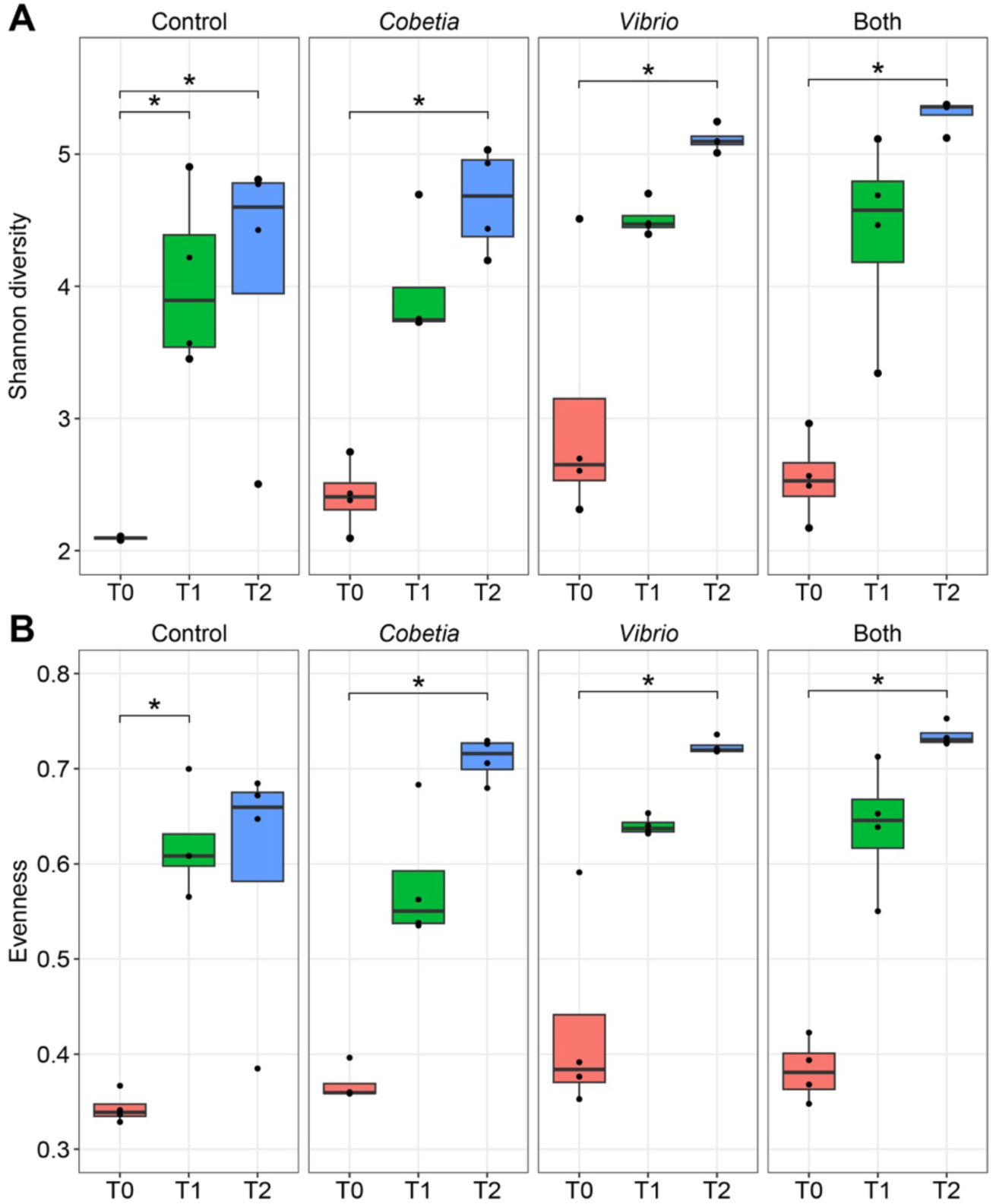
Alpha diversity indices from the microbiomes of the heat-stressed samples, separated by bacteria inoculation and timepoint. The y-axes represent (A) the alpha diversity measured using Shannon diversity and (B) the evenness within each sample, measured using the Pielou index. Samples were compared across timepoints within each inoculation group using Kruskal Wallis H tests, followed by Dunn’s multiple comparison tests (n = 4 for each group). Significant differences (p < 0.05, after Benjamini-Hochberg correction) are denoted with asterisks.

**Extended Data Fig. 3.**
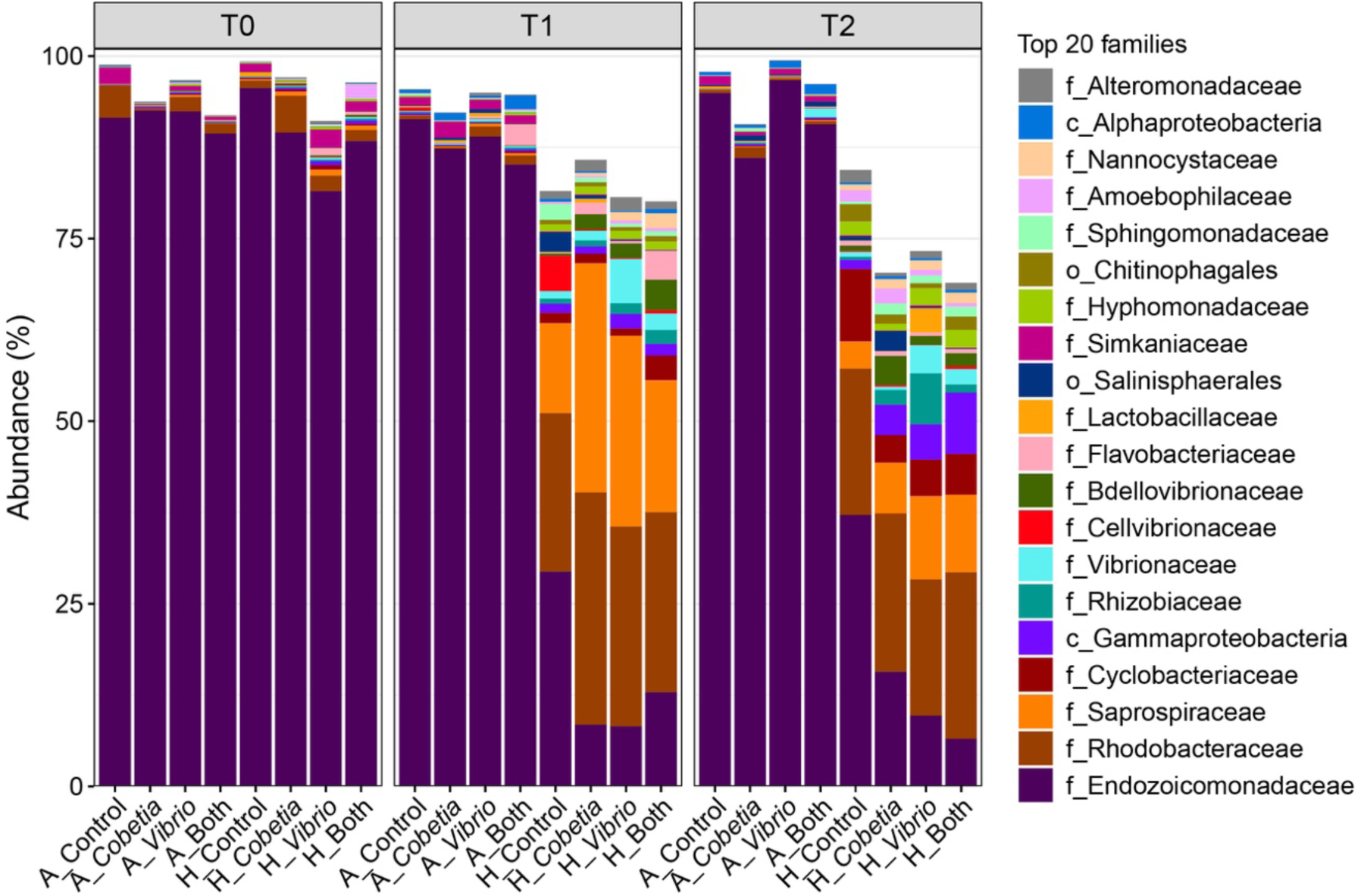
The top 20 most abundant bacterial families within the samples are represented as stacked barcharts for each experimental group, separated by timepoint (n = 4 per group). The y-axis is the relative abundance for each bacterium within the sample group. For the bacteria, where the family was not identified, the lowest taxonomic identification is listed (“f_” = family, “c_” = class, “o_” = order).

**Extended Data Fig. 4.**
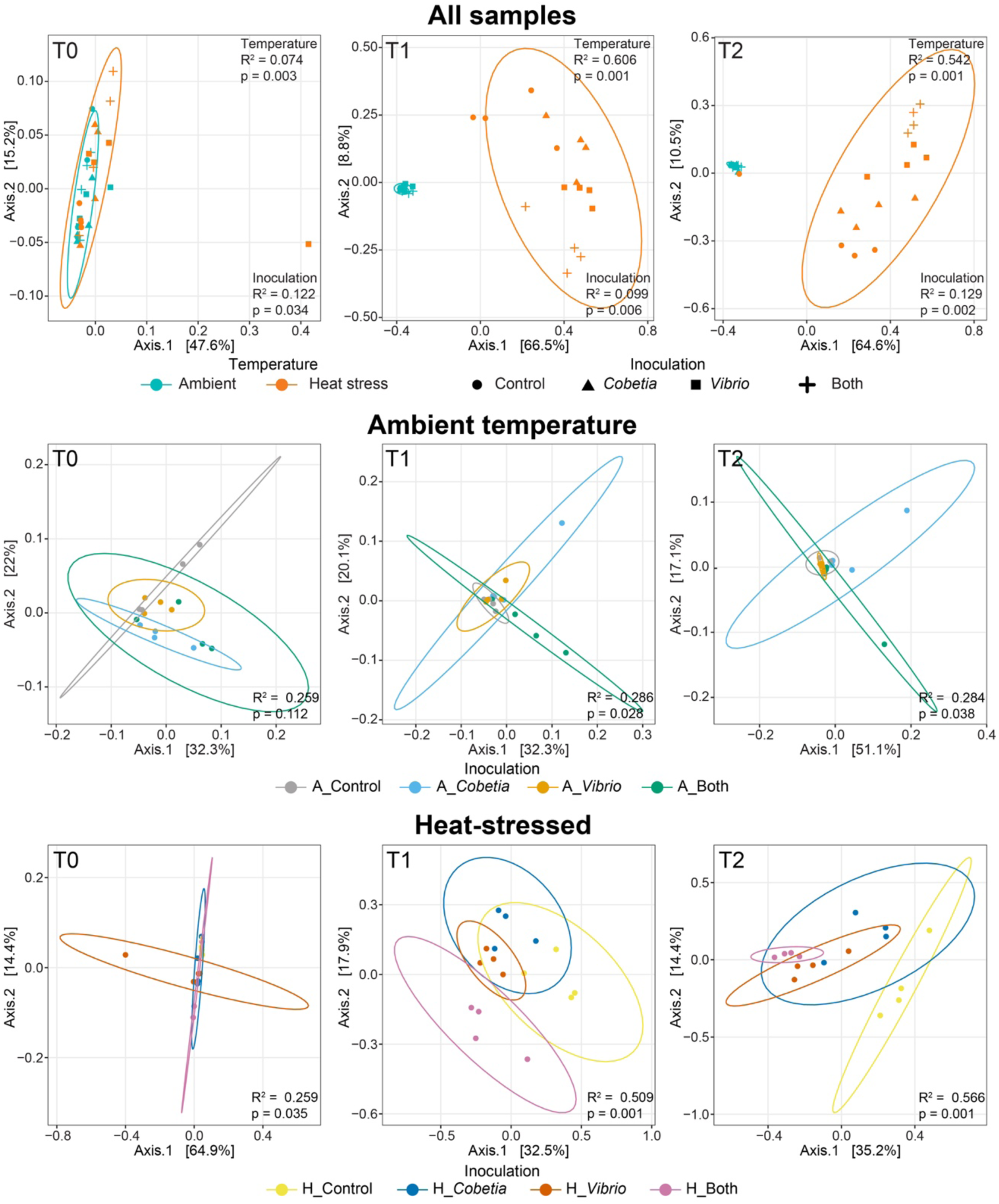
Principle coordinate analysis (PCoA) plots based on Bray-Curtis dissimilarities of the relative abundances of the total bacterial community in the coral nubbins at each sampling time. Samples from each timepoint were separated using Bray-Curtis dissimilarity and plotted along the first two dimensions (n = 4 per group). Circles = 95% confidence intervals around temperature groups. Both temperature and inoculation were tested as factors driving the separation between groups using PERMANOVA (top row). For each temperature regime (ambient – middle row and heat-stressed – bottom row), significant differences between inoculation were tested using PERMANOVA. R^2^ and p-values are displayed, representing the magnitude and significance attributed to each of the factors.

**Extended Data Fig. 5.**
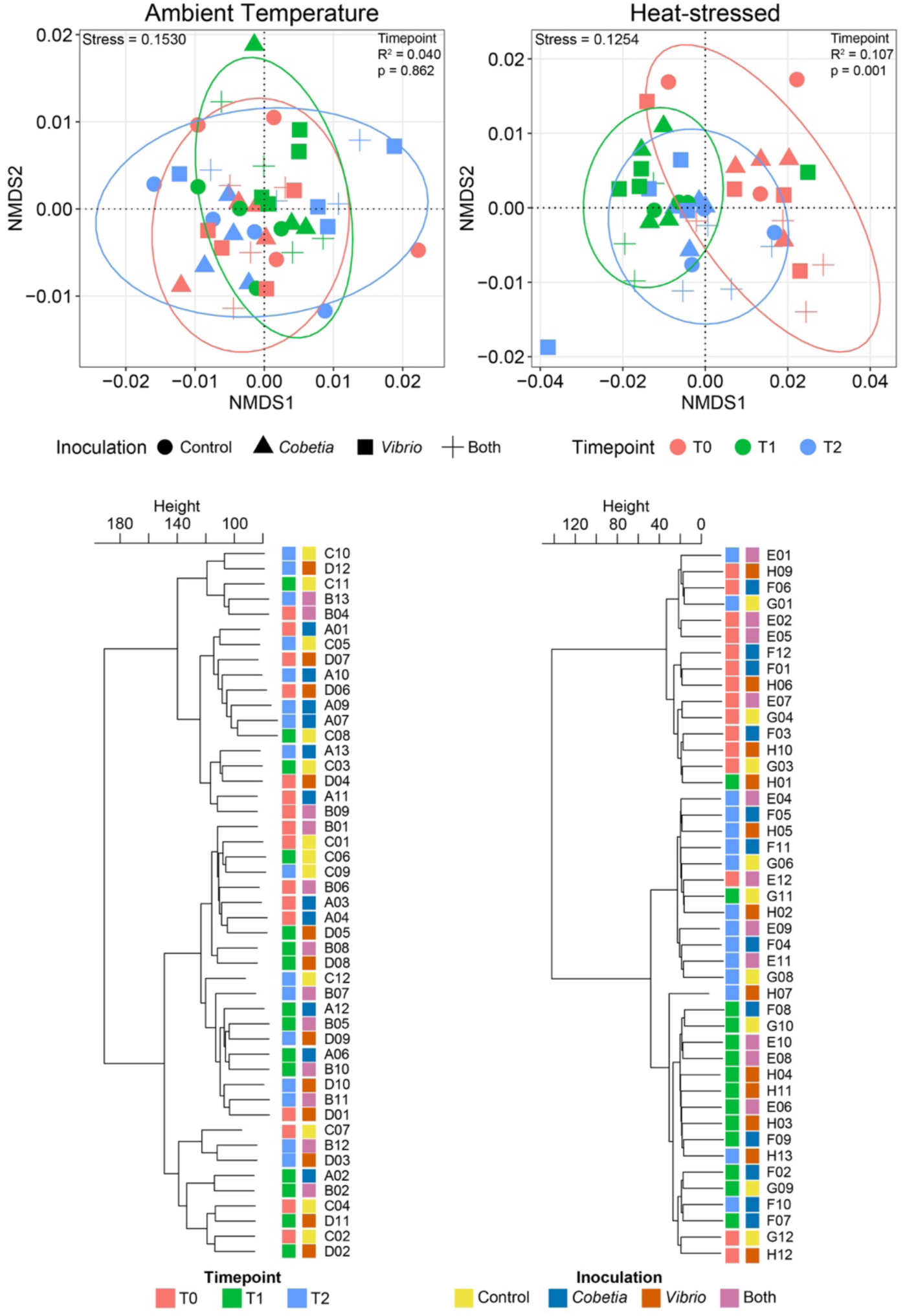
Separation of genome-wide DNA methylation in samples based on timepoint in ambient temperature corals and heat-stressed corals via nMDS plots using Bray-Curtis dissimilarity and hierarchical clustering using the “ward.D2” agglomeration method. Circles in nMDS plots = 95% confidence intervals around timepoints. For each temperature regime (ambient – left and heat-stressed – right), significant differences between timepoints were tested using PERMANOVA (Ambient Temperature, T0/T2: n = 16 and T1: n = 15; Heat-stressed, T0/T2: n = 15 and T1: n = 14). R^2^ and p-values are displayed, representing the magnitude and significance attributed to each of the factors. Hierarchical clustering demonstrates the relative dissimilarity between clusters of samples based on methylation levels across the genome (represented by dendrogram height).

**Extended Data Fig. 6.**
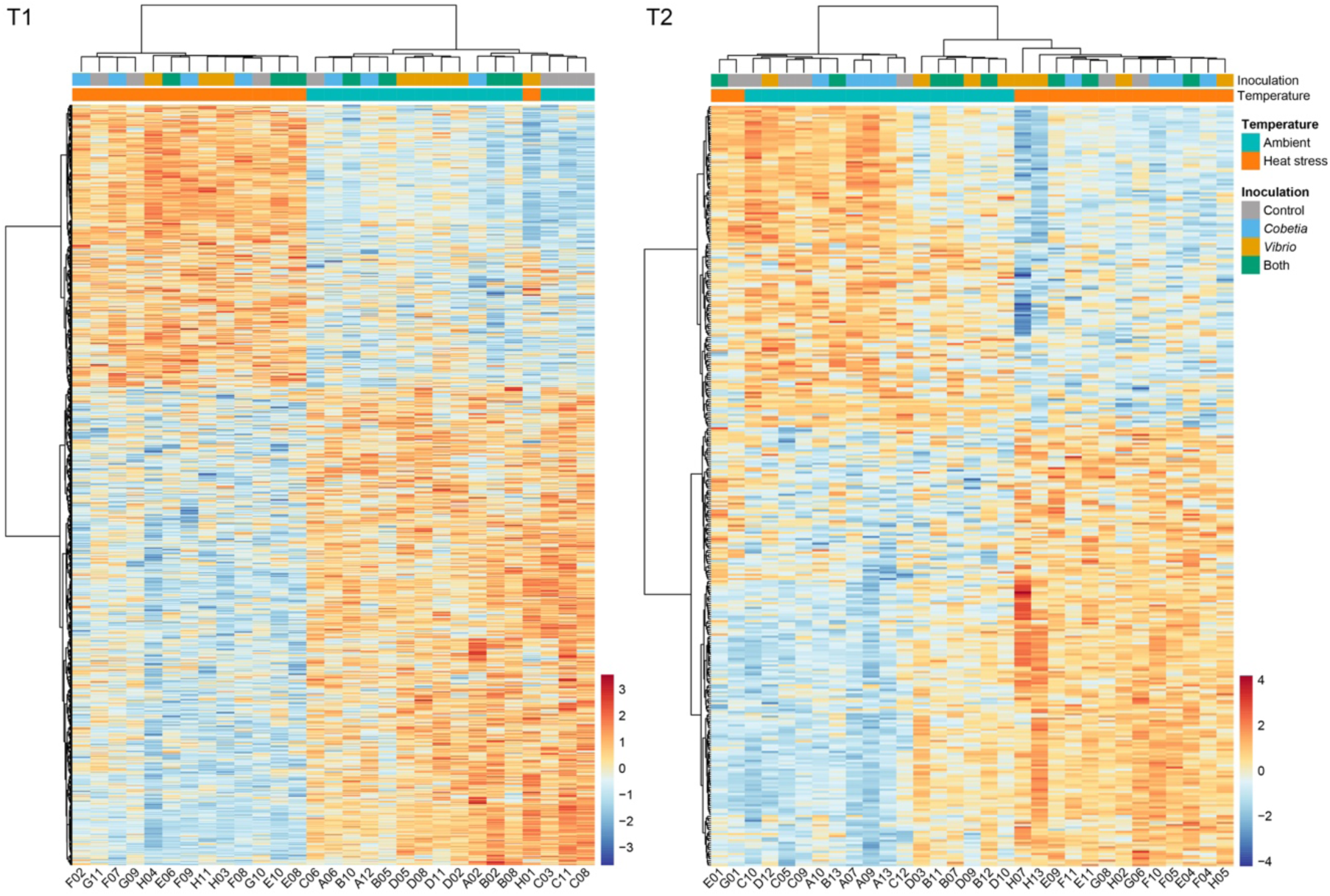
Percent methylation levels of genes significantly driven by temperature conditions (ambient vs. heat stress) at sampling times T1 (left) and T2 (right) as determined by GLMs (n = 4; n = 3 for H_Control at T1/T2; n = 3 for H_Both at T1). Hierarchical clustering was performed using Euclidean distances and the “ward.D2” agglomeration method. Colors represent the z-score for any given gene. All enriched GO terms due to differences in temperature regimes at T1 and T2 are in Supplementary Table 6a and 6b, respectively.

**Extended Data Fig. 7.**
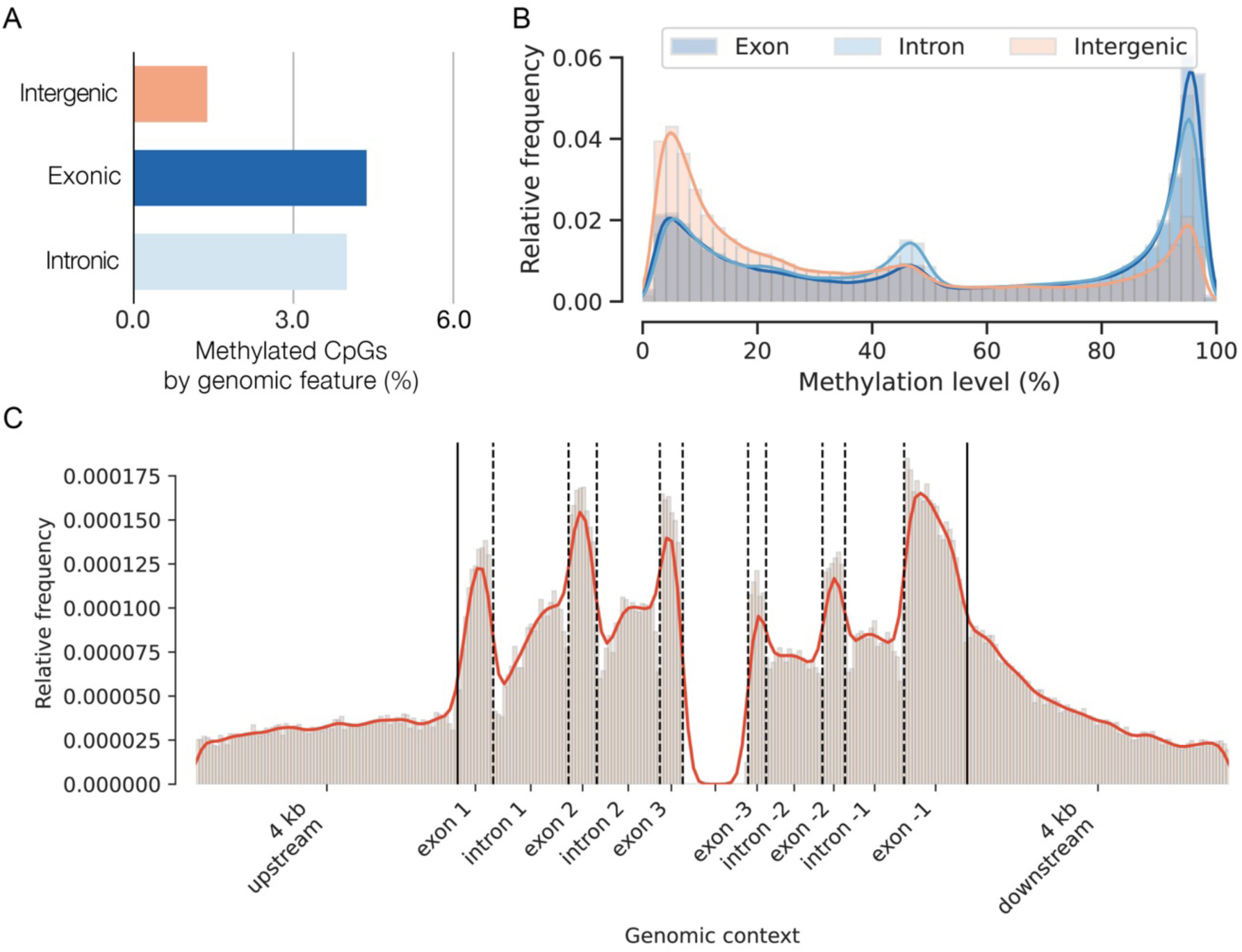
Genomic context of methylation across genomic regions and along genes. Data from all samples (n = 91) were pooled to find where DNA methylation is observed along the *Pocillopora verrucosa* genome. (A) Percent methylated CpGs in each genomic feature group. (B) Methylation levels display approximate bimodal distribution, with higher methylation levels in genic regions (exons and introns) than intergenic regions. (C) The relative frequencies of methylated positions along a standardized gene model show than methylated positions are more frequent within exons.

**Extended Data Fig. 8.**
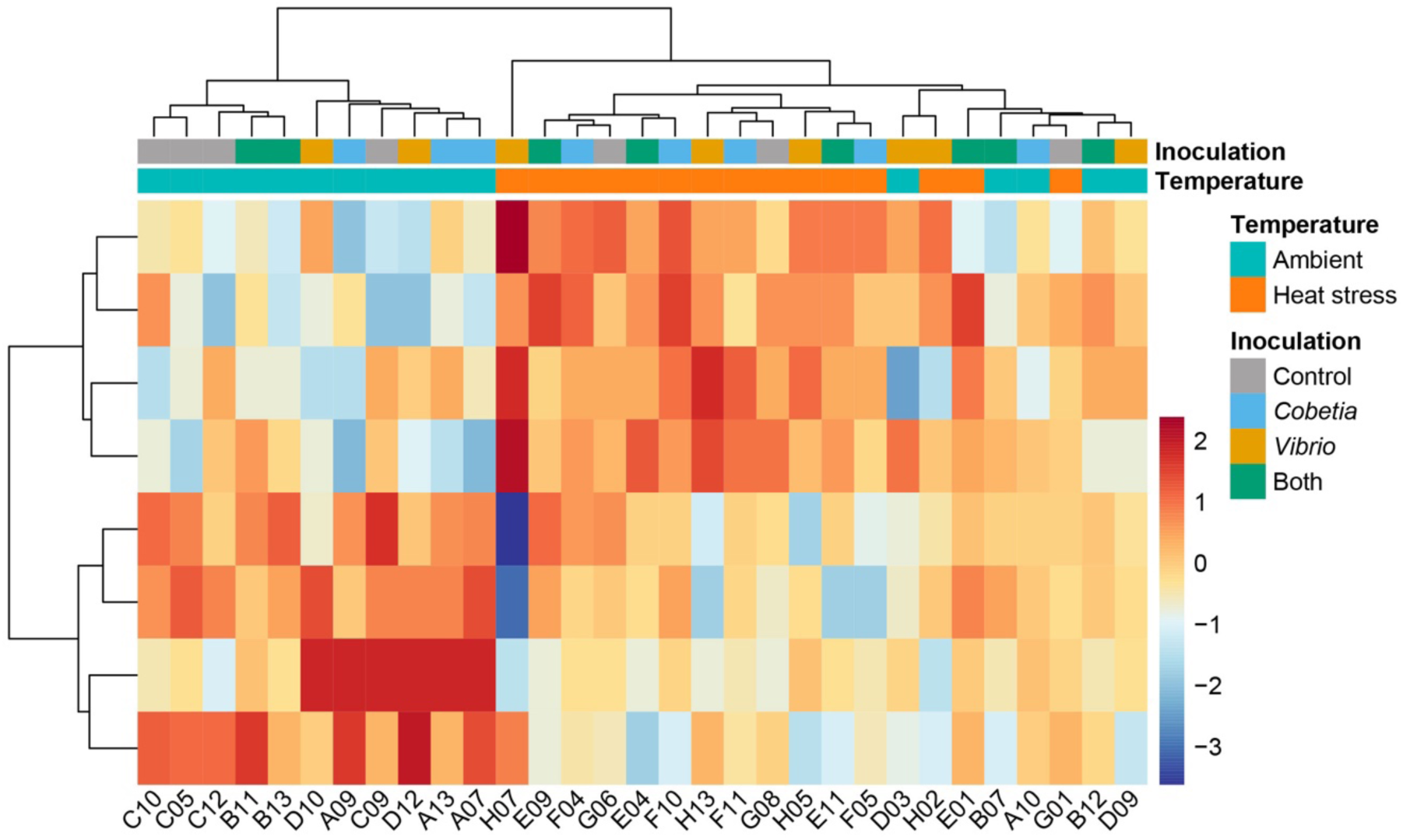
Percent methylation levels of glutamatergic synapse genes significantly driven by temperature conditions (ambient vs. heat stress) at T2 (8 genes total) as determined by GLMs (n = 4; n = 3 for H_Control). Hierarchical clustering was performed using Euclidean distances and the “ward.D2” agglomeration method. Colors represent the z-score for any given gene.

**Extended Data Fig. 9.**
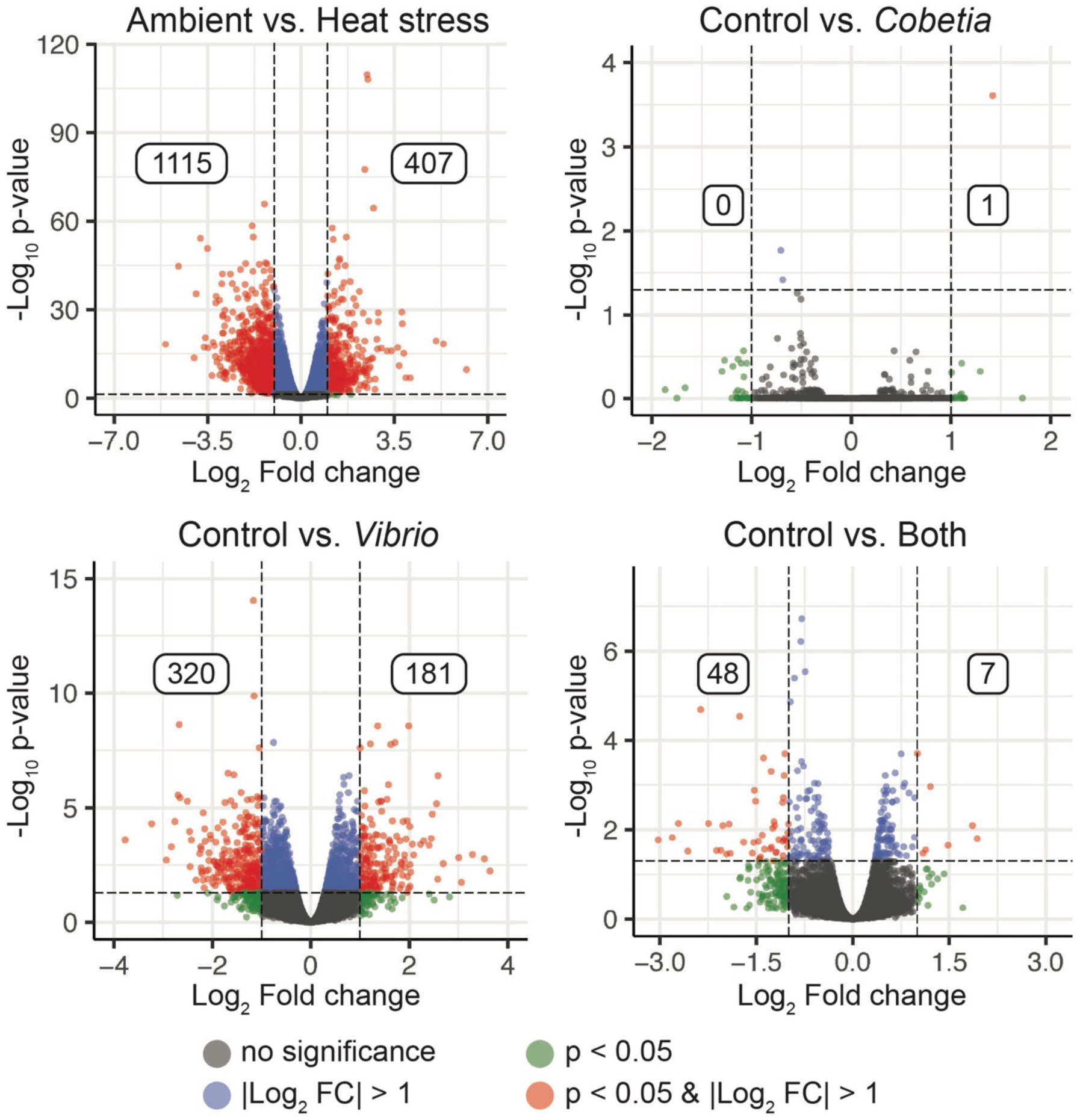
Volcano plots displaying fold-change comparisons from RNA-seq data between the temperature conditions (ambient vs. heat stress; n = 16) and each of the inoculation groups to the control group (n = 8). Each dot within the volcano plots represents the normalized count of a gene. Significantly up-regulated or down-regulated genes in comparison to the ambient temperature (in the temperature comparison) or the control group (in the inoculation comparisons) was determined by a log_2_ fold change >1 or <-1 and a p < 0.05 after adjusting for false discovery rate.

## Notes

### Competing Interest Statement

The authors have declared no competing interest.

### Summary of Updates

The text was revised and new RNA-seq data was added.

## References

1. Doering, T. et al. Towards enhancing coral heat tolerance: a “microbiome transplantation” treatment using inoculations of homogenized coral tissues. Microbiome 9, 102 (2021)

2. Fragoso ados Santos, H., et al. Impact of oil spills on coral reefs can be reduced by bioremediation using probiotic microbiota. Sci. Rep. 5, 18268 (2015)

3. Moradi, M. et al. Probiotics mitigate thermal stress- and pathogen-driven impacts on coral skeleton. Front. Mar. Sci. 10, 1212690 (2023)

4. Rosado, P. M. et al. Marine probiotics: increasing coral resistance to bleaching through microbiome manipulation. ISME J. 13, 921–936 (2019)

5. Santoro, E. P. et al. Coral microbiome manipulation elicits metabolic and genetic restructuring to mitigate heat stress and evade mortality. Sci. Adv. 7, eabg3088 (2021)

6. Silva, D. P. et al. Multi-domain probiotic consortium as an alternative to chemical remediation of oil spills at coral reefs and adjacent sites. Microbiome 9, 118 (2021)

7. Ushijima, B. et al. Chemical and genomic characterization of a potential probiotic treatment for stony coral tissue loss disease. Commun. Biol. 6, 248 (2023)

8. Peixoto, R. S., Rosado, P. M., Leite, D. C. de A., Rosado, A. S. & Bourne, D. G. Beneficial Microorganisms for Corals (BMC): Proposed Mechanisms for Coral Health and Resilience. Front. Microbiol. 8, 341 (2017)

9. Reshef, L., Koren, O., Loya, Y., Zilber-Rosenberg, I. & Rosenberg, E. The Coral Probiotic Hypothesis. Environ. Microbiol. 8, 2068–2073 (2006)

10. Rohwer, F., Seguritan, V., Azam, F. & Knowlton, N. Diversity and distribution of coral-associated bacteria. Mar. Ecol. Prog. Ser. 243, 1–10 (2002)

11. Zhang, Y. et al. Shifting the microbiome of a coral holobiont and improving host physiology by inoculation with a potentially beneficial bacterial consortium. BMC Microbiol. 21, 130 (2021)

12. Sweet, M. J. & Bulling, M. T. On the Importance of the Microbiome and Pathobiome in Coral Health and Disease. Front. Mar. Sci. 4, 9 (2017)

13. Barno, A. R., Villela, H. D. M., Aranda, M., Thomas, T. & Peixoto, R. S. Host under epigenetic control: A novel perspective on the interaction between microorganisms and corals. BioEssays 43, 10 (2021)

14. Li, Y. et al. DNA methylation regulates transcriptional homeostasis of algal endosymbiosis in the coral model Aiptasia. Sci. Adv. 4, 8 (2018)

15. Liew, Y. J. et al. Epigenome-associated phenotypic acclimatization to ocean acidification in a reef-building coral. Sci. Adv. 4, 6 (2018)

16. Neri, F. et al. Intragenic DNA methylation prevents spurious transcription initiation. Nature 543, 72–77 (2017)

17. Rodriguez-Casariego, J. A., Cunning, R., Baker, A. C. & Eirin-Lopez, J. M. Symbiont shuffling induces differential DNA methylation responses to thermal stress in the coral *Montastraea cavernosa*. Mol. Ecol. 31, 588–602 (2021)

18. Putnam, H. M. Avenues of reef-building coral acclimatization in response to rapid environmental change. J. Exp. Biol. 224, jeb239319 (2021)

19. Raimundo, I., Rosado, P. M., Barno, A. R., Antony, C. P. & Peixoto, R. S. Unlocking the genomic potential of Red Sea coral probiotics. Sci. Rep. 14, 14514 (2024)

20. Ben-Haim, Y. et al. *Vibrio coralliilyticus* sp. nov., a temperature-dependent pathogen of the coral *Pocillopora damicornis*. Int. J. Syst. Evol. Microbiol. 53, 309–315 (2003)

21. Siebeck, U. E., Logan, D. & Marshall, N. J. CoralWatch – a flexible coral bleaching monitoring tool for you and your group. Coral Reefs. (2010)

22. Cui, G. et al. Host-dependent nitrogen recycling as a mechanism of symbiont control in *Aiptasia*. PLOS Genet. 15, e1008189 (2019)

23. Rädecker, N. et al. Heat stress destabilizes symbiotic nutrient cycling in corals. Proc. Natl. Acad. Sci. 118, e2022653118 (2021)

24. Vega Thurber, R., et al. Deciphering Coral Disease Dynamics: Integrating Host, Microbiome, and the Changing Environment. Front. Ecol. Evol. 8, 575927 (2020)

25. Dimond, J. L. & Roberts, S. B. Convergence of DNA Methylation Profiles of the Reef Coral *Porites astreoides* in a Novel Environment. Front. Mar. Sci. 6, 792 (2020)

26. Dixon, G., Liao, Y., Bay, L. K. & Matz, M. V. Role of gene body methylation in acclimatization and adaptation in a basal metazoan. Proc. Natl. Acad. Sci. 115, 13342–13346 (2018)

27. Dixon, G. B., Bay, L. K. & Matz, M. V. Bimodal signatures of germline methylation are linked with gene expression plasticity in the coral *Acropora millepora*. BMC Genomics 15, 1109 (2014)

28. Putnam, H. M., Davidson, J. M. & Gates, R. D. Ocean acidification influences host DNA methylation and phenotypic plasticity in environmentally susceptible corals. Evol. Appl. 9, 1165–1178 (2016)

29. Liew, Y. J. et al. Intergenerational epigenetic inheritance in reef-building corals. Nat. Clim. Change 10, 254–259 (2020)

30. Rodríguez-Casariego, J. A., et al. Coral epigenetic responses to nutrient stress: Histone H2A.X phosphorylation dynamics and DNA methylation in the staghorn coral *Acropora cervicornis*. Ecol. Evol. 8, 12193–12207 (2018)

31. Guerrero, L. & Bay, R. Patterns of methylation and transcriptional plasticity during thermal acclimation in a reef-building coral. Evol. Appl. 17, e13757 (2024)

32. Abbott, E., Loockerman, C., & Matz, M. V. Modifications to gene body methylation do not alter gene expression plasticity in a reef-building coral. Evol. Appl. 17, e13662 (2024)

33. Rodríguez-Casariego, J. A. et al. Genome-Wide DNA Methylation Analysis Reveals a Conserved Epigenetic Response to Seasonal Environmental Variation in the Staghorn Coral *Acropora cervicornis*. Front. Mar. Sci. 7, 560424 (2020)

34. Turgut-Kara, N., Arikan, B. & Celik, H. Epigenetic memory and priming in plants. Genetica 148, 47–54 (2020)

35. Konciute, M. K. Exploring the Role of Glutamate Signaling in the Regulation of the Aiptasia-Symbiodiniaceae Symbiosis. KAUST Research Repository (2020)

36. Kawamura, K. et al. *In vitro* Symbiosis of Reef-Building Coral Cells With Photosynthetic Dinoflagellates. Front. Mar. Sci. 8, 706308 (2021)

37. Kamran, Z. et al. In vivo imaging of epithelial wound healing in the cnidarian *Clytia hemisphaerica* demonstrates early evolution of purse string and cell crawling closure mechanisms. BMC Dev. Biol. 17, 17 (2017)

38. Agarwal, S. et al. Autophagy and endosomal trafficking inhibition by *Vibrio cholerae* MARTX toxin phosphatidylinositol-3-phosphate-specific phospholipase A1 activity. Nat. Commun. 6, 8745 (2015)

39. Kimes, N. E. et al. Temperature regulation of virulence factors in the pathogen *Vibrio coralliilyticus*. ISME J. 6, 835–846 (2012)

40. Dixon, G. & Matz, M. V. Changes in gene body methylation do not correlate with changes in gene expression in Anthozoa or Hexapoda. BMC Genomics 23, 234 (2022)

41. Lehnert, E. M., Mouchka, M. E., Burriesci, M. S., Gallo, N. D., Schwarz, J. A., & Pringle, J. R. Extensive Differences in Gene Expression Between Symbiotic and Aposymbiotic Cnidarians. G3 Genes|Genomes|Genetics 4, 277–295 (2014)

42. Pacherres, C. O., Ahmerkamp, S., Koren, K., Richter, C., & Holtappels, M. Ciliary flows in corals ventilate target areas of high photosynthetic oxygen production. Curr. Biol. 32, 4150–4158 (2022)

43. Hansen, S. B. et al. Intestinal epigenotype of Atlantic salmon (*Salmo salar*) associates with tenacibaculosis and gut microbiota composition. Genomics 115, 110629 (2023)

44. Garcias-Bonet, N. et al. Horizon scanning the application of probiotics for wildlife. Trends Microbiol. S0966-842X(23)00259-7 (2023)

45. Peixoto, R. S., Sweet, M. & Bourne, D. G. Customized Medicine for Corals. Front. Mar. Sci. 6, 686 (2019)

46. Peixoto, R. S. & Voolstra, C. R. The baseline is already shifted: marine microbiome restoration and rehabilitation as essential tools to mitigate ecosystem decline. Front. Mar. Sci. 10, 1218531 (2023)

47. Voolstra, C. R. & Ziegler, M. Adapting with microbial help: microbiome flexibility facilitates rapid responses to environmental change. BioEssays. 42, 7 (2020)

48. Dixon, G. & Matz, M. Benchmarking DNA methylation assays in a reef-building coral. Mol. Ecol. Resour. 21, 464–477 (2021)

49. Wilcoxon, F. Individual Comparisons by Ranking Methods. Biometrics Bulletin 1, 80–83 (1945)

50. Kruskal, W. H. & Wallis, W. A. Use of ranks in one-criterion variance analysis. J. Am. Stat. Assoc. 47, 583–621 (1952)

51. Dunn, O. J. Multiple Comparisons Using Rank Sums. Technometrics 6, 241–252 (1964)

52. Martin, M. Cutadapt removes adapter sequences from high-throughput sequencing reads. EMBnet.journal 17, 10–12 (2011)

53. Callahan, B. J. et al. DADA2: High-resolution sample inference from Illumina amplicon data. Nat. Methods 13, 581–583 (2016)

54. Quast, C. et al. The SILVA ribosomal RNA gene database project: improved data processing and web-based tools. Nucleic Acids Res. 41, D590–D596 (2013)

55. McMurdie, P. J. & Holmes, S. phyloseq: An R Package for Reproducible Interactive Analysis and Graphics of Microbiome Census Data. PLoS ONE 8, e61217 (2013)

56. Altschul, S. F., Gish, W., Miller, W., Myers, E. W. & Lipman, D. J. Basic Local Alignment Search Tool. J Mol Biol 215, 403–410 (1990)

57. Buitrago-López, C., Mariappan, K. G., Cárdenas, A., Gegner, H. M. & Voolstra, C. R. The Genome of the Cauliflower Coral *Pocillopora verrucosa*. Genome Biol. Evol. 12, 1911–1917 (2020)

58. Krueger, F. & Andrews, S. R. Bismark: a flexible aligner and methylation caller for Bisulfite-Seq applications. Bioinformatics 27, 1571–1572 (2011)

59. Alexa, A., Rahnenführer, J. & Lengauer, T. Improved scoring of functional groups from gene expression data by decorrelating GO graph structure. Bioinformatics 22, 1600–1607 (2006)

60. Bray, N. L., Pimentel, H., Melsted, P. & Pachter, L. Near-optimal probabilistic RNA-seq quantification. Nat. Biotechnol. 34, 525–527 (2016)

61. Pimentel, H. J., Bray, N., Puente, S., Melsted, P. & Pachter, L. Differential analysis of RNA-seq incorporating quantification uncertainty. Nat. Methods 14, 687–690 (2017)

